# Single-cell copy number calling and event history reconstruction

**DOI:** 10.1101/2020.04.28.065755

**Authors:** Jack Kuipers, Mustafa Anıl Tuncel, Pedro F. Ferreira, Katharina Jahn, Niko Beerenwinkel

## Abstract

Copy number alterations are driving forces of tumour development and the emergence of intra-tumour heterogeneity. A comprehensive picture of these genomic aberrations is therefore essential for the development of personalised and precise cancer diagnostics and therapies. Single-cell sequencing offers the highest resolution for copy number profiling down to the level of individual cells. Recent high-throughput protocols allow for the processing of hundreds of cells through shallow whole-genome DNA sequencing. The resulting low read-depth data poses substantial statistical and computational challenges to the identification of copy number alterations. We developed SCICoNE, a statistical model and MCMC algorithm tailored to single-cell copy number profiling from shallow whole-genome DNA sequencing data. SCICoNE reconstructs the history of copy number events in the tumour and uses these evolutionary relationships to identify the copy number profiles of the individual cells. We show the accuracy of this approach in evaluations on simulated data and demonstrate its practicability in applications to two breast cancer samples from different sequencing protocols.

## Introduction

During tumour progression, cancer cells undergo complex and diverse genomic aberrations leading to heterogeneous cell populations and multiple evolving subclones [1, 2, 3]. Genomic sequencing has been powerfully employed to uncover the heterogeneity across cancer types [4] and to examine the link between tumour diversity and progression [5, 6, 7]. Heterogeneity may also allow the tumour additional ways to evolve resistance under treatment, such that intra-tumour genomic diversity is a cause of relapse and treatment failure [8, 9].

The need for a comprehensive understanding of the composition of each tumour for more precise and effective cancer therapies [10] can be addressed by sequencing tumours at the resolution of individual cells [11, 12]. The small amount of DNA in single cells has to be amplified before sequencing, which leads to characteristic and pronounced noise in the read count data. Specialised phylogenetic methods for single-cell sequencing data accounting for these noise patterns have been developed to detect point mutations and reconstruct the evolutionary history of tumours [13, 14].

In addition to point mutations, cancerous cells often undergo more complex genomic rearrangements, including copy number alterations (CNAs) such as amplifications and deletions. Since the first single-cell DNA sequencing [15], which allowed for the identification of CNAs, rapid progress has led to high-throughput methods [16, 17] that can profile the whole genome of hundreds of single cells. By removing the need for pre-amplification, the protocols of [16, 17] have particularly uniform coverage allowing a resolution of CNAs down to the megabase scale. A commercial solution from 10x Genomics was also available for a time, likewise enabling the processing of hundreds of single cells, and employed in a large scale clinical project [18]. The resolution and depth of the data depends heavily on the technology employed, and the amount of sequencing subsequently performed.

A number of computational tools have been applied or specifically developed for copy number calling in single-cell DNA sequencing data [19, 20, 21, 22, 23, 24, 25] with reviews and comparisons in an overview paper [26]. The output of these methods is a copy number profile, a partition of the genome into segments and a sequence of integers indicating the copy number state of each segment. While some of these tools pool information across cells [23, 24], the tools above do not take into account the shared evolutionary history of cells originating from the same individual or tumour. As previously shown for point mutations, the evolutionary relationships of tumour cells can be used to boost and correct the weak and noisy signal provided by single-cell sequencing reads [27]. However, modelling and inference of evolutionary histories from single-cell data is notably more involved in the case of CNAs, as such events may physically overlap, and will more readily reoccur and revert.

Methods for jointly calling copy numbers and reconstructing event histories are therefore currently under active development. Previous methods developed for inferring evolutionary histories based on copy numbers assume that copy number profiles of the individual cells are predefined. The methods can be separated into two categories, distance-based approaches [15, 28, 29]) and methods that reconstruct the evolutionary history based on a maximum parsimony heuristic [30, 31, 32, 33]. Further distinction can be made by the way identical copy number changes of neighbouring segments are interpreted, as separate events [34], a single event [28, 30, 29], or a hybrid thereof in which deletions/amplifications can either occur separately on individual segments or affect whole chromosomes or genomes [31, 32]. A limitation shared by all these methods is that errors in the copy number profiles will propagate into the inference of the copy-number trees.

Our approach, SCICoNE (https://github.com/cbg-ethz/SCICoNE), directly integrates the inference of copy number profiles with the reconstruction of copy number event histories tailored to the shallow read-depth of whole-genome single-cell sequencing data. Other tree-based methods have also been developed [35, 36]: CONET [35] builds the tree based on the likelihood of the presence or absence of break-points and calls copy-numbers in post-processing using a discrepancy penalty although this does not directly model the evolution of copy-numbers, while NestedBD [36] works in the cell lineage space. We therefore benchmark SCICoNE against a large suite of alternatives, including tree-based methods, and show strong improvements in performance, especially in the low read-depth and high noise regime typical of very shallow whole-genome single-cell sequencing, used for example in [18]. We further employ SCICoNE to examine the copy number history of a xenograft breast cancer sample, and a breast cancer tissue sample processed with the 10x Genomics platform.

## Results

### Model overview

During tumour evolution, CNAs can occur and accumulate in cancer cell sub-populations. Since all cells in a tumour are related through sharing a common ancestor, CNAs occur on a cell lineage tree and encode the copy number profiles of each clone or cell (Figure 1a). With the whole-genome sequencing of single cells [16, 37], we have read depth data for each cell and each bin along the genome (Figure 1b). After correction for confounders, including read mappability and genomic GC content, for each bin [38, 20], the corrected counts can be assumed to be proportional to the underlying copy number. To partition the genome into segments of consecutive bins which may have experienced CNAs, we develop a dynamic programming approach to detect breakpoints by combining evidence across the individual cells (Methods). For each potential breakpoint, we compare the likelihood of a step change in copy number state to that of a constant model, and then collate the information across cells to obtain the total signal of a copy number change at that genomic position.

**Figure 1:**
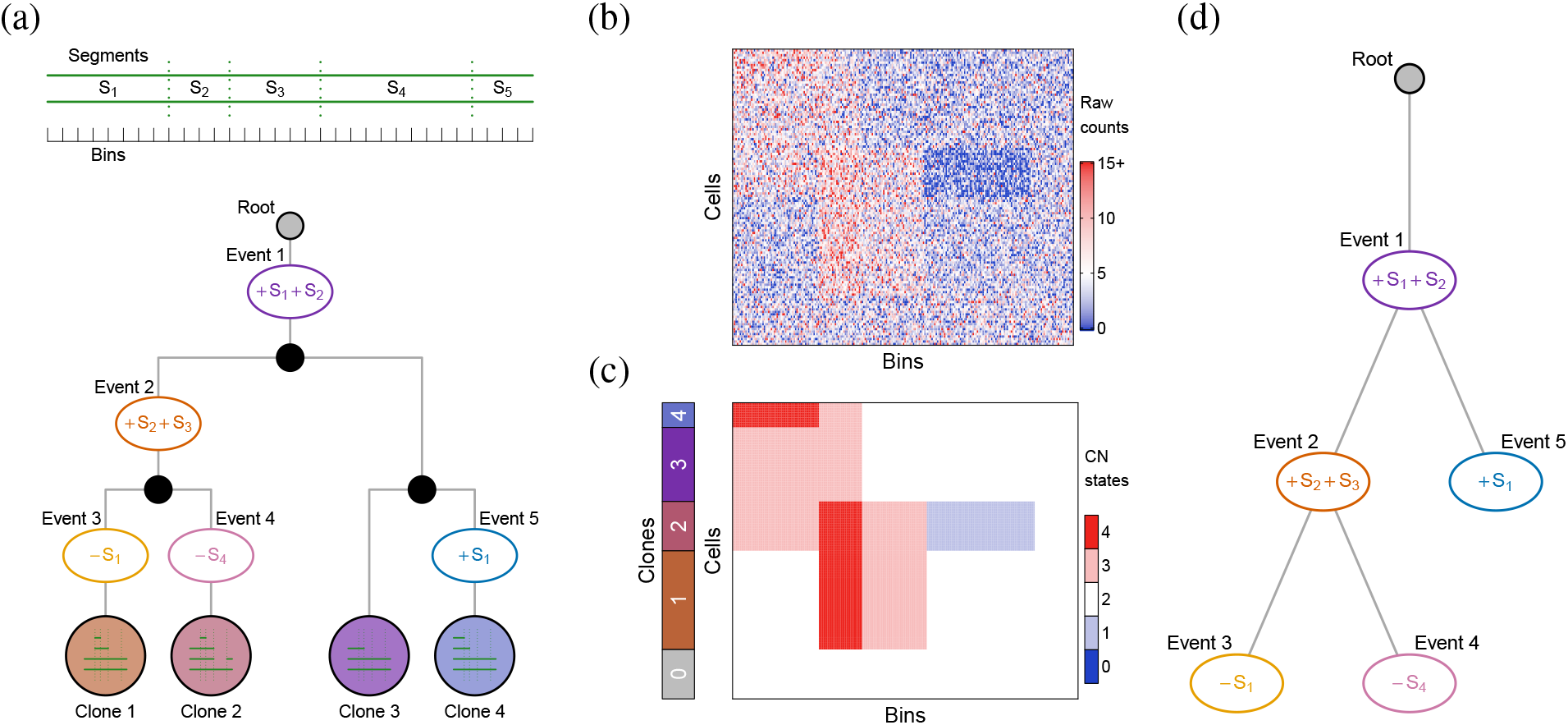
CNA calling and tree inference. (a) The genome is partitioned, by the four breakpoints depicted, into the five segments *S*_1_, …, *S*_5_ (each comprising several bins) that may experience CNAs. During tumour evolution, copy number changes may accumulate along the branches (plus signifies an amplification, minus a deletion) of the cell lineage tree to lead to distinct tumour subclones. The copy number profile of each clone can be obtained by tracing the lineages from the root, or neutral diploid cells. For example, Clone 4 has experienced two CN events, namely Event 1, a gain in a region spanning segments *S*_1_ and *S*_2_, and Event 5, a further gain in *S*_1_, such that the copy number profile of Clone 4 is (4, 3, 2, 2, 2) across the five segments. (b) From single-cell sequencing we obtain noisy read count data according to the copy number profiles of the clones and their sizes. (c, d) Accounting for lineage relationships between cells and modelling the noise in the data, SCICoNE infers both the copy number calls in each cell (c) and the sequence of the CNAs and their phylogenetic relationships (d).

For clinical applications, the history of copy number events is often more pertinent than the fully resolved cell genealogy, as the former is sufficient to determine which CNAs are mutually exclusive or co-occurring in the same tumour subclone, and to infer the order of CNA events. We therefore move from the cell or clone lineage tree (Figure 1a) to the CNA tree (Figure 1d). To account for the noise in the sequencing data, we develop a probabilistic model and an MCMC inference scheme for single-cell read counts (Methods). A major difference to MCMC schemes used for reconstructing trees of point mutations [39] is that the infinite sites assumption, which excludes the possibility of multiple mutational hits at the same genomic site, can no longer be made [40]. CNAs may overlap and nest inside each other, so the model developed here allows for arbitrary violations of the infinite sites assumption and arbitrary reoccurrences of amplifications and deletions across different genomic regions. Reoccurrences are only resolved and modelled at the level of the bins of the genomic data, so that the individual break points may still occur at different genomic positions in the same bin. Along with allowing violations of the infinite sites assumption at the bin level, we explicitly model dependencies between bins as they are tied together within copy number events according to the CNA tree model.

By jointly estimating the copy number profiles of all cells and their underlying CNA tree, we leverage the evolutionary dependencies among cells for improved copy number calls. The output of SCICoNE comprises both (1) the reconstructed tree representing the evolutionary history of all cells (Figure 1d) and (2) the inferred copy number profile for each individual cell (Figure 1c).

### Reconstructing copy number trees from tumour data

As a first demonstration of SCICoNE on whole-genome single-cell sequencing data, we considered a dataset of 260 cells of a triple negative breast cancer xenografted into a mouse (SA501X3F) [16]. The segmentation algorithm of SCICoNE (Methods, with default parameters) revealed 316 candidate breakpoints over the 18,175 bins.

The reconstructed tree (Figure 2 and Supplementary Figure A1) starts with a large number of CNAs and then branches into two main populations which further differentiate into different clones. The first node includes deletions in *TP53* and the *MAPK* genes along with an amplification in *PIK3CA*. Notably, *AKT1* undergoes repeated amplifications, while *ARID1B* and *ESR1* are first amplified at the root and later undergo a deletion in one sub-branch. The genes *NTRK3* and *TBX3* instead experience deletions each in two parallel lineages. Overall, the tree highlights the complex evolutionary history of the sample.

**Figure 2:**
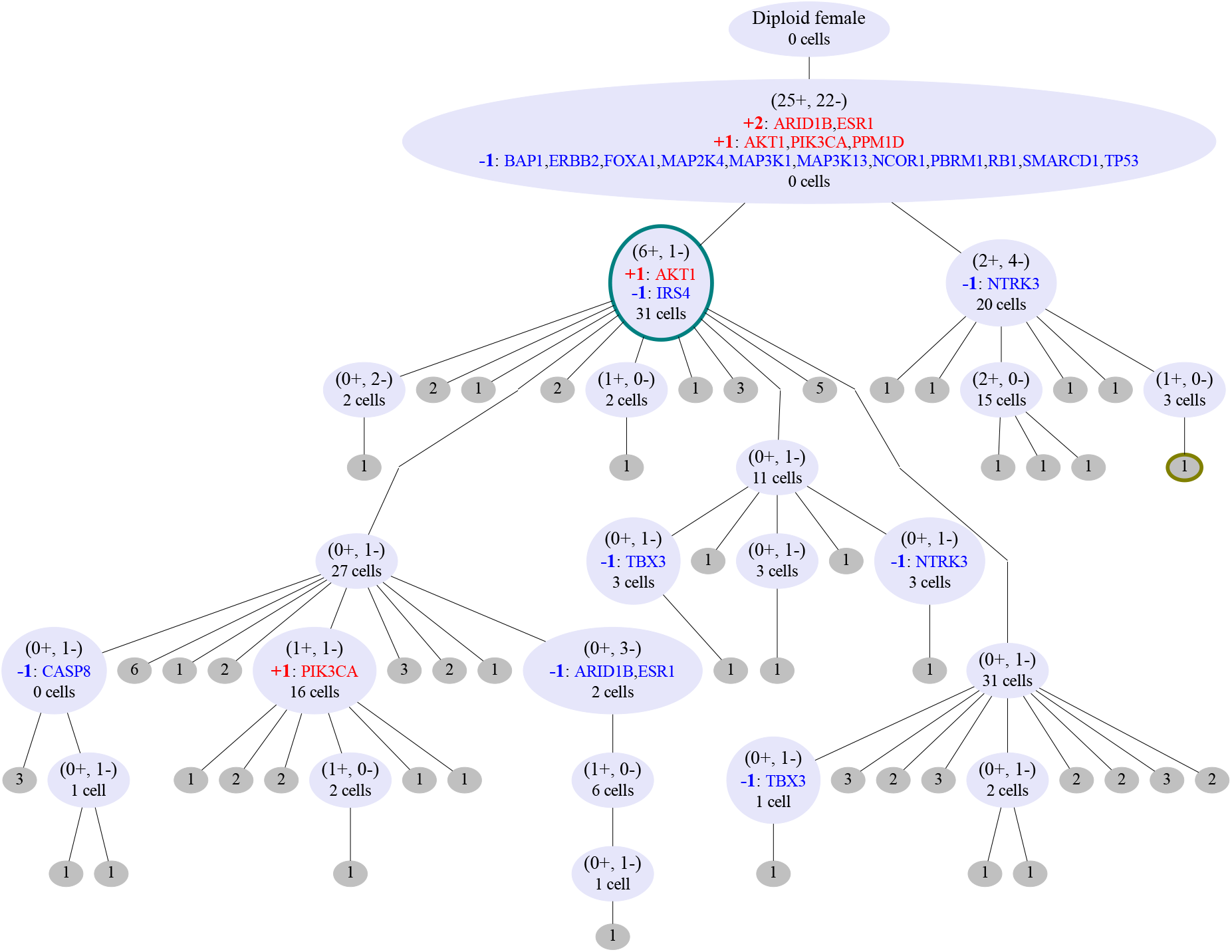
Inferred tree for 260 cells from a breast xenograft [16]. Inside the nodes of the CNA tree we highlight the total number of amplification or deletion events (in parentheses), the genes which are affected (among the 33 associated with breast cancer in the COSMIC Cancer Gene Census [41]), including how much they are amplified or deleted, and the number of cells that best attach to each node. The CNAs are not displayed at the grey (leaf) nodes where only the number of cells attached is indicated. Example profiles of cells attaching to the two nodes with coloured borders are displayed in Figure 3c.

The inferred copy number profiles (Figure 3b) are in good agreement with the normalised counts per bin (Figure 3a), recapitulating the CNAs across cells while accounting for the noise in the raw data and the phylogenetic relationships between the cells. Joint inference of copy number profiles and their evolutionary history provides increased power to separate signal from noise, as emphasised by comparing the raw count and inferred copy number profiles of example cells (Figure 3c). While some small changes for individual cells visible in the normalised counts (Figure 3a) are filtered out, most changes occurring even in a small number of cells are detected (Figure 3b). Overall, we obtain the main CNAs across cells as well as their phylogeny (Figure 2).

**Figure 3:**
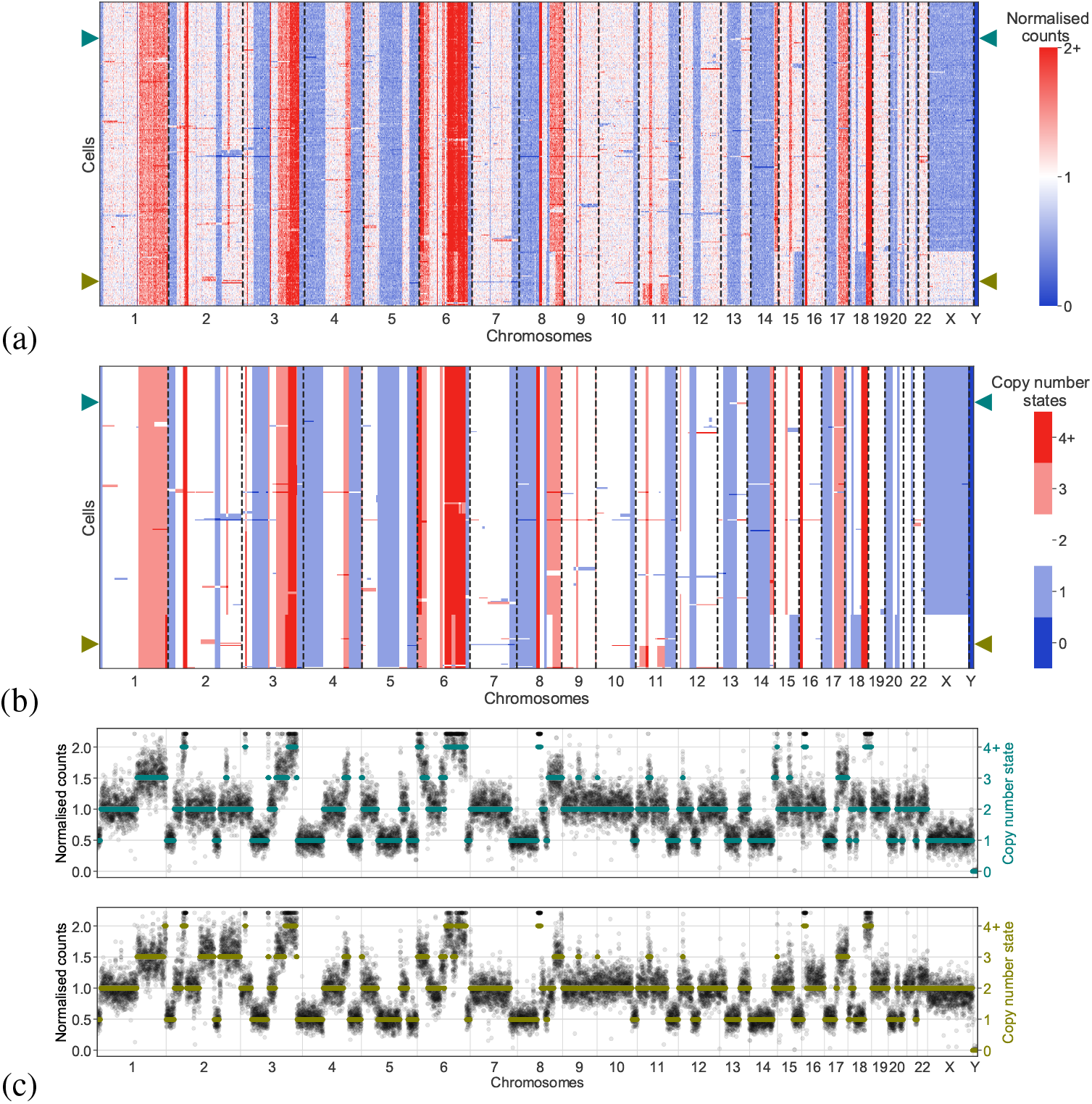
Inferred copy number profiles for 260 cells from a breast xenograft [16]. (a) Normalised counts per bin, ordered according to the tree in Figure 2. (b) Copy number profiles estimated jointly with the CNA tree of Figure 2. (c) Two examples of raw count data (black dots) and inferred copy number profiles (coloured lines) of the two cells indicated by arrows in the heatmaps

To examine the relationship between CNAs and RNA expression changes at the single-cell level, we analysed the xenograft passage SA501X2B which underwent single-cell RNA sequencing [42] and compared the two molecular profiles. By smoothing the RNA signal [43], we observe common strips of over- and underexpression (Supplementary Figure A2). Comparing to the DNA summarised at the gene level (by averaging over the bins in each of the 6000 expressed genes), we find some agreement between genomic copy number changes and expression levels (Supplementary Figure A2). However, quite a few of the signals visible in the RNA data, like in chromosome 19, for example, have no basis in the DNA. The concordance and discrepancies between RNA expression and DNA count levels is further emphasised if we cluster the cells (Supplementary Figure A3). The finer structure and clear breakpoints visible in the DNA, which are reflected in the inferred evolutionary history (Figure 2) and copy number profiles (Figure 3) are not well reflected in the RNA (Supplementary Figure A2). Moreover the correlation between RNA expression profiles and DNA read counts, while present, is fairly weak (Supplementary Figure A4). The results confirm on the level of individual cells that expression profiles are not in a strict one-to-one correspondence with copy number profiles and that the latter can be inferred with higher accuracy and in more detail from DNA data.

Next we examined 2053 single-cells from a triple negative breast cancer available as an example dataset from 10x Genomics [37] (Breast Tissue nuclei section E). The 10x Genomics Cellranger pipeline filtered out 45 cells as low quality and performed GC correction, while we removed 57 outlier bins with more than 3 times the median counts. To robustly detect breakpoints from the 20kb-sized bins, we used a window size of 100 bins, allowing for the detection of copy number events that span at least 2Mb. Already from the read counts per cell (Figure 4b), we can see a difference between the clusters based on the read-count profiles (Figure 4c), with higher levels in the tumour cells (clones 1-3) pointing to a possible whole-genome duplication. Comparing the results of SCICoNE with a diploid root against a run with a tetraploid root account for a genome duplication, we observe much higher likelihoods in the latter case. The resulting cluster tree and copy number calls (Figure 4a and d) recapitulate the main clonal architecture and their evolutionary relationships.

**Figure 4:**
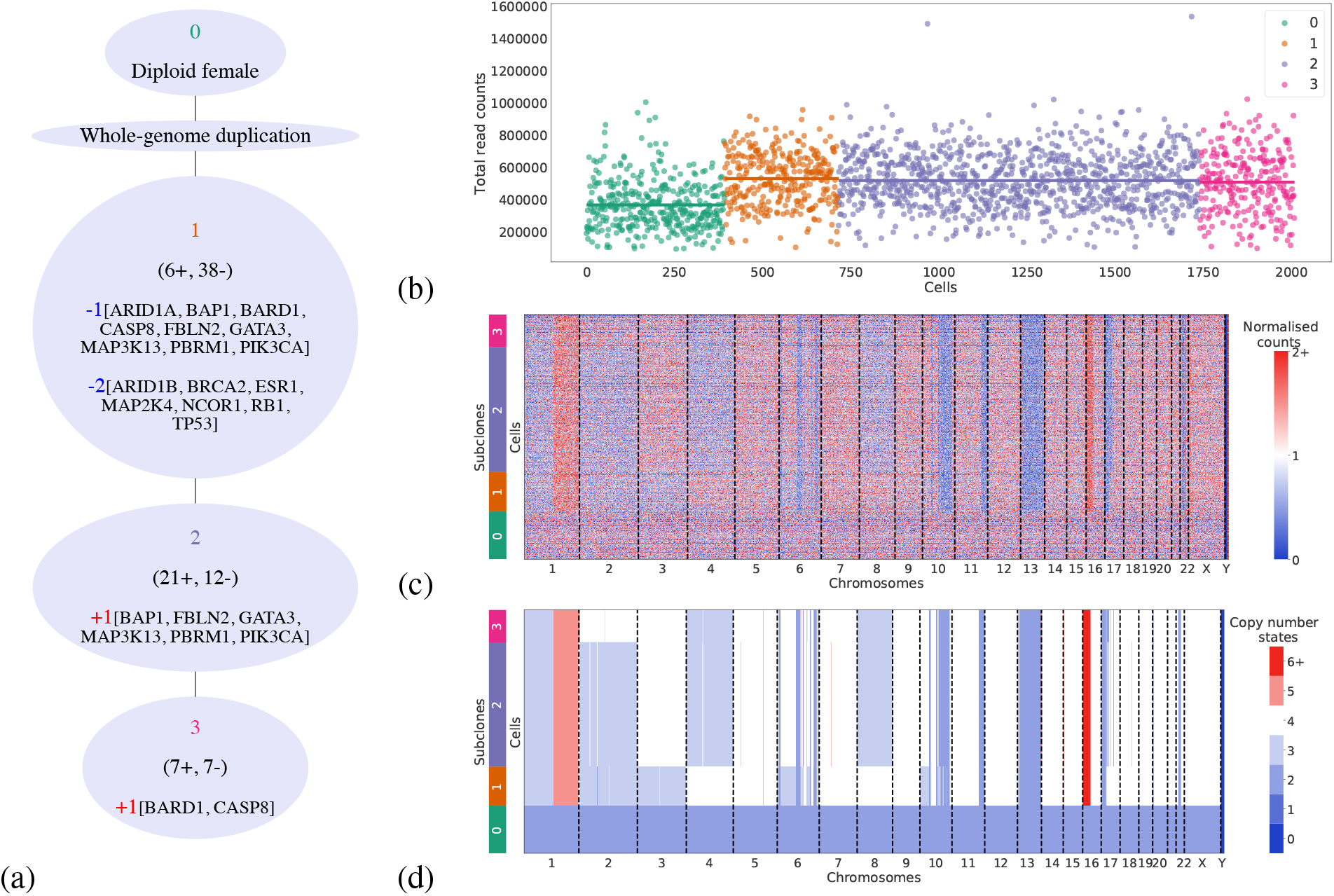
Inferred copy number profiles for 2053 cells from a breast cancer [37]. (a) Inferred tree on the clustered data, with the genes which are affected (among the 33 associated with breast cancer in the COSMIC Cancer Gene Census [41]) displayed at each node. (b) Counts per cell for the different clusters. (c) Normalised counts per bin. (d) Copy number profiles estimated jointly with the CNA tree.

### Benchmarking on simulated data

To benchmark SCICoNE we conducted a simulation study (Methods), mimicking the very shallow coverage of a recent large clinical project [18], and compared the performance of SCICoNE to a suite of alternatives. For simulated trees with 20 nodes (Figure 5), we observe a steady increase in accuracy as we model our data in more detail. We first consider methods that cluster the data, and as a baseline, we cluster the normalised count data using hierarchical clustering or PhenoGraph [44] (Figure 5, gold and silver) and assign cells the averaged profile of their clusters (rounded to integers and centred at diploid). Then we build trees on PhenoGraph clustered data using SCICoNE to leverage information across the clusters and their shared evolutionary history to improve the accuracy of learning the copy number profiles (Figure 5, red and pink). Next we compare to methods that work at the single-cell level, and we perform the full tree inference with SCICoNE (Figure 5, purple and blue), observing a strong improvement over the alternatives. On the full data, we find that the ‘max’ setting, which finds the best placement of each cell in the CNA tree, offers higher reconstruction quality relative to ‘sum’, which averages over their placements (Methods, Equation (9) and Equation (10)). We therefore used ‘max’ for the breast cancer data above.

**Figure 5:**
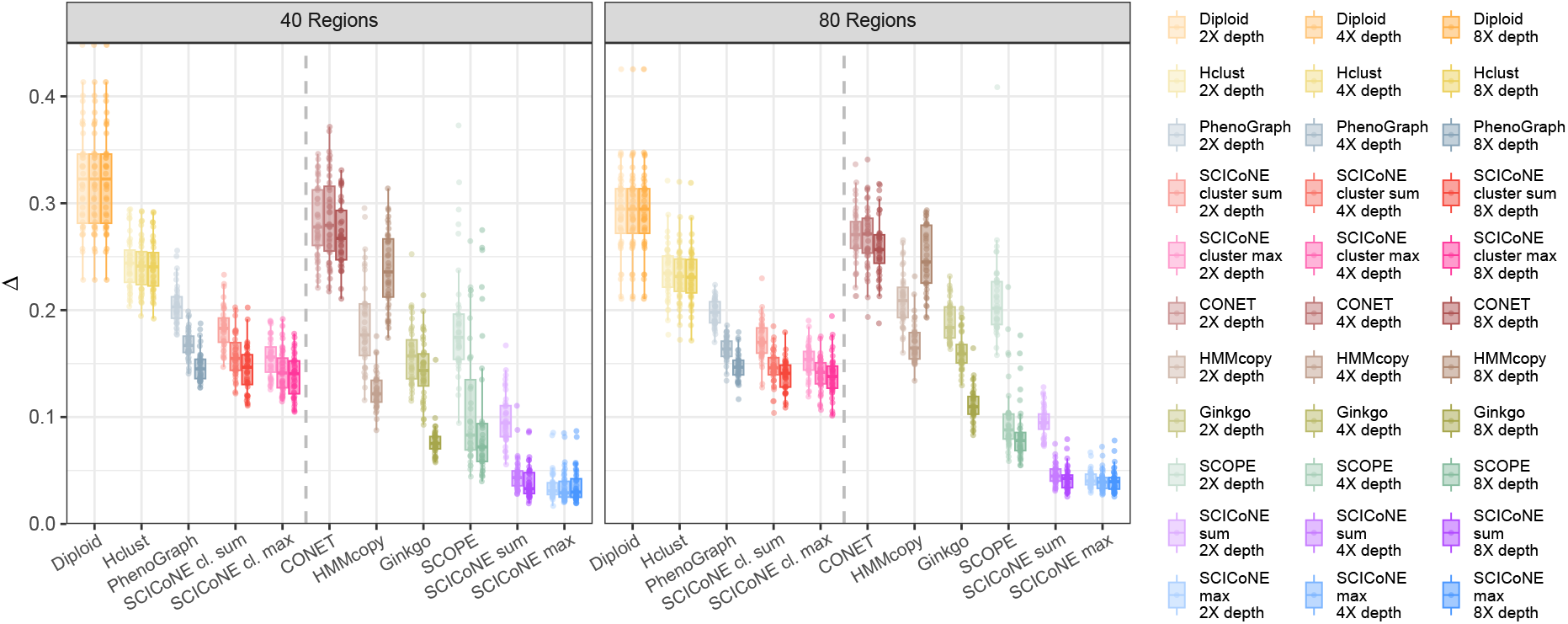
Comparison of copy number calling for simulated data. For random trees with 20 nodes, we attached 400 cells and simulated overdispersed read data according to each cell’s copy number profile over 10,000 bins. The total number of reads was 20k, 40k and 80k for an average read depth of 2X, 4X and 8X (colour intensity, left to right) per bin for each cell. The maximal number of segments affected by copy number changes was 40 and 80 (panels). The root mean squared difference Δ between the true simulated copy number profiles and the corresponding inferred profiles over all bins and cells is summarised in each box plot (generated with ggplot2 default settings), for a neutral diploid profile (orange), for profiles inferred by hierarchical (gold) and PhenoGraph (silver) clustering as well as SCICoNE on PhenoGraph clustered data (red and pink), followed by CONET [35] (reddish brown) and HMMcopy [19] (bronze), Ginkgo [20] (olive) and SCOPE [23] (green), and SCICoNE on the full single-cell sequencing data (purple and blue).

SCICoNE performs well in comparison to HMMcopy [19] (Figure 5, bronze) which performs copy number calling per cell with a hidden Markov model, and has been found to have good overall performance in single-cell copy number calling [26]. However, for the simulated data with an average read depth of 2–8 (comparable to 10x Genomics data), HMMcopy can have quite variable performance and is generally worse than the Phenograph clustering (Figure 5, silver). Instead we found Ginkgo [20] (Figure 5, olive) to perform similarly to PhenoGraph clustering at lower depths and with more segments (and smaller copy number events) but better with fewer segments and especially at the highest depth. SCOPE [23] (Figure 5, green) is better still at the higher depths and a little worse than Ginkgo at the lowest depth, although its performance is quite variable across the repetitions. In contrast, SCICoNE on the clustered data is very robust to the read depth and performs better than the other clustering approaches (hierarchical and PhenoGraph), while on the full (un-clustered) data SCICoNE allows us to extract much more accurate copy number profiles than alternative (HMMcopy, Ginkgo and SCOPE) methods, along with the evolutionary history encoded in the tree itself. We observe similar performances in the easier setting of higher coverage (more akin to the coverage of the protocols of [16, 17]) and no overdispersion (Supplementary Figure A5), although HMMcopy performs notably worse due to it often calling the reference level incorrectly, while using SCICoNE to build a tree on the clustered data does not offer an advantage in copy number calling compared to the clustering itself.

CONET [35], despite using a phylogenetic tree, performs very poorly in the simulations, at odds with the results they report. We tracked this discrepancy down to their simulations implicitly providing CONET with effectively much higher read-depth data than they provided to SCICoNE, and perform a detailed analysis in Appendix E. To benchmark the tree learning we take the output of Ginkgo and SCOPE as input for MEDALT [33] which reconstructs a phylogenetic tree from called copy numbers, and compare to the trees from CONET [35] and SCICoNE (Figure 6). Similarly to the copy number calling (Figure 5), CONET is the worst performer, but close to the performance of Ginkgo with MEDALT. SCOPE offers a better input to MEDALT than Ginkgo at the higher depths, but worse at the lowest, with quite high variability as a function of depth. Overall the best performance comes from modelling the data with SCICoNE especially with the ‘max’ setting, in line with the results for copy number calling.

**Figure 6:**
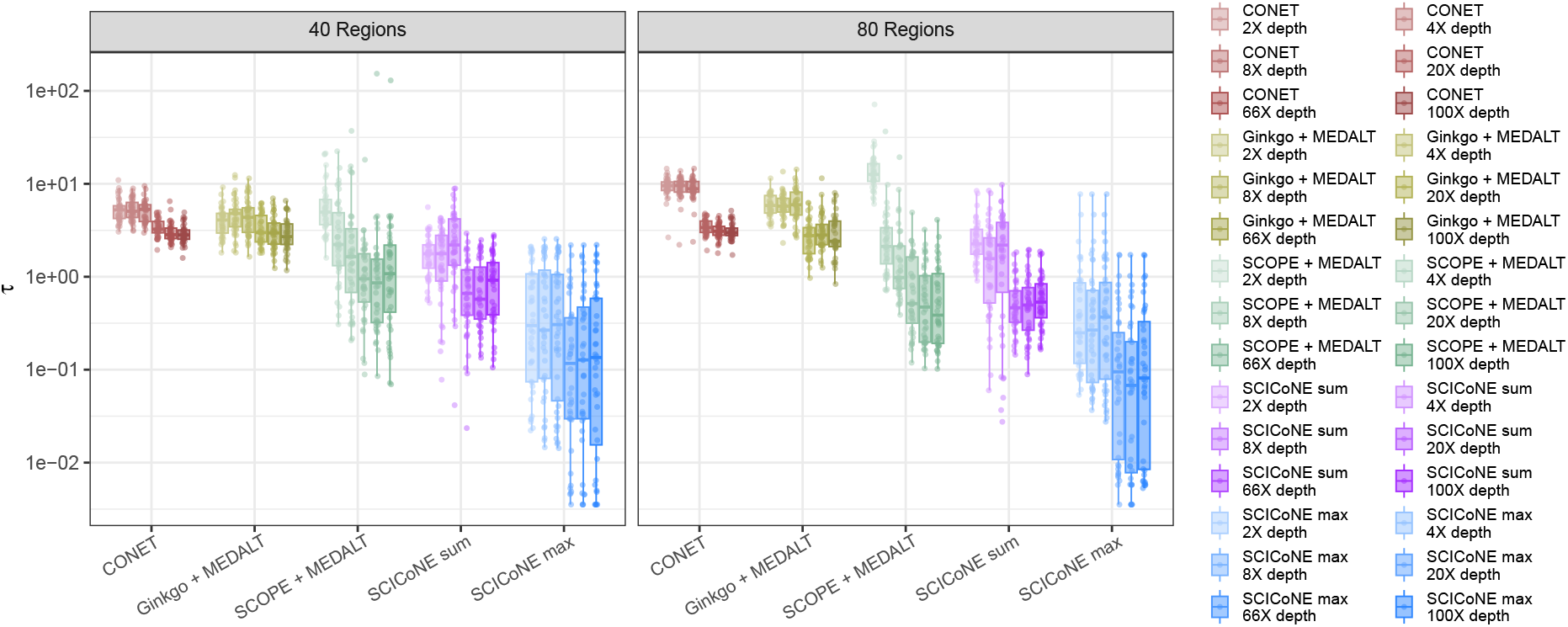
Comparison of copy number tree reconstruction for simulated data. For the simulated data of Figure 5 and Supplementary Figure A5 we compute the tree distance *τ* (Methods) between the true and inferred tree from CONET [35] (reddish brown), MEDALT [33] run on the output of Ginkgo [20] (olive) and SCOPE [23] (green), as well as SCICoNE (purple and blue).

## Conclusions

For learning the copy number profiles of single cells, sharing information across cells and leveraging their shared evolutionary history boosts our ability to remove noise and accurately call copy numbers. Here we developed a novel phylogenetic framework for this purpose, enabling us to jointly infer the tree and copy number states and obtain better quality profiles. When learning the sequence of CNAs that occur in tumour samples from single-cell sequencing data, the possible overlap and reoccurrences of CNAs need to be accounted for. Our tree model allows regions of the genome to be arbitrarily amplified and deleted, while controlling for model complexity. Simulations demonstrate that our approach is accurate for the challenging task of reconstructing the evolutionary history of tumours and calling copy numbers in individual cells.

For a real data example of 260 xenograft breast cancer cells, we obtained the CNA tree and inferred copy number profiles of the cells and detected the main clonal structure as well as their phylogenetic relationship. Some smaller or weaker changes over small numbers of cells, visible in the normalised data (Figure 3b) were not detected as copy number changes in the inference (Figure 3a) as they were insufficient to justify a more complex model in our framework. Adjusting and relaxing the penalisation for model complexity to detect finer changes may offer further improvements, though learning larger and more complex trees also increase the computation cost of the inference.

For the 2053 cells from a triple negative breast ductal carcinoma, sequenced with the 10x Genomics technology, we could detect a whole-genome amplification and use this to correct the ploidy for the tumour cells to accurately call their copy number states.

To learn the segments in the first place, we developed a dynamic programming approach to combine the evidence of breakpoints across all cells. Compared to methods which work on a per cell basis, our joint inference of breakpoints and then the full probabilistic model of the phylogeny provides a substantial improvement. The combination of bins into segments reduces the possible search space and speeds up the inference, but since the quality of the segments detected directly affects the downstream reconstruction, further improvements in this direction are important. In particular, once a phylogeny has been learnt based on the strongest breakpoints, the corresponding separation of cells into clones can help distinguish noise from signal in the breakpoint detection. Adding and removing breakpoints could also be incorporated as a move into the MCMC scheme itself.

For real data, where GC and mappability corrections are performed to normalise the bins, residual confounding effects still remain which can complicate the breakpoint detection and phylogenetic inference. In particular, the bin corrections depend on the underlying copy number state, indicating that future directions could consider jointly inferring the corrections along with the segments and the phylogeny, although increasing the complexity and computational cost of learning CNA trees.

Understanding and reconstructing the history of copy number events in a tumour could play a key role in predicting response to treatment, especially when resistance arises from adaptive selection of the existing clonal architecture. The combination with single-cell transcriptomics can uncover the interplay between evolutionary pressure and cellular reprogramming [45]. The phylogenetic methods developed here for large-scale, complex and overlapping events could potentially also reconstruct trajectory structures reflected in the transcriptomic profiles of single-cell RNA sequencing, while the copy number trees reflect a common structure on which to analyses expression profiles [42, 46].

Although developed for shallow whole-genome single-cell sequencing protocols [16, 17, 18], our phylogenetic methods may also play a key role in copy number reconstruction for targetted single-cell sequencing [47], and offer relevant methodology for phylogenetics which integrate copy numbers and point mutations [48] and for considering allele-specific events where methods currently look at the phasing through mutations but without jointly learning the phylogenetic history [24, 49]. Copy number reconstruction further complements multi-faceted single-cell profiling [18], for example to determine the downstream effects of tumour heterogeneity through evolutionary analyses across cohorts. Accurate copy number calling at the single-cell level, enabled through SCICoNE’s joint inference of the evolutionary history and copy number profiles, will enhance single-cell analysis for cancer biology.

## Methods

### Binning and read count correction

Current protocols for copy number detection at single-cell resolution typically employ shallow whole-genome sequencing (≲0.1x coverage per cell) [16, 37] which prohibits coverage-based copy number calling at the level of individual loci. Instead one partitions the reference genome into equal sized bins (20–100kb) and counts the reads per bin instead of per locus. The raw read count of a bin, is not only determined by the bin’s underlying copy number state, but also by its mappability and GC content. To reduce the bias introduced by these confounders, SCICoNE uses read counts (per bin and per cell) that have been corrected for both effects [38, 20]. With these confounders removed, we now assume that the probability of reads falling into each bin is proportional to the bin’s copy number state,

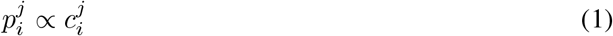

Where 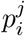 is the probability of a read from cell *j* falling into bin *i*, and 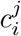 is the copy number state for that bin in that cell.

### Breakpoint detection to define copy number segments

As copy number changes often affect regions much larger than a typical bin size, we collate neighbouring bins into genomic segments with the same copy number state. To collate the bins into genomic segments, we first detect potential break-points as bin boundaries where the read depth changes across subsets of cells. For this purpose we developed a dynamic programming approach that combines evidence across cells to call the breakpoints (Appendix A). The detection compares a likelihood-based model of a step change in copy number at each bin to a constant copy number model for each cell and then combines the signal over all cells. Bins with the strongest combined signal relative to a noise threshold are classified as potential breakpoints.

Once the breakpoints have been determined, we collate bins between consecutive breakpoints into segments. For each cell, we sum the counts in all bins belonging to each segment to arrive at a count matrix *D* with entries *D*_*jk*_ for each cell *j* and each segment *S*_*k*_.

The probability of a read falling into segment *S*_*k*_ is proportional to the copy number state and the size of the segment

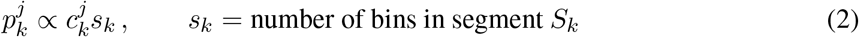

Where 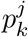 is the probability of a read from cell *j* falling into segment *k* with size *s*_*k*_, and 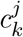 is the copy number state for that segment in that cell.

### CNA trees

For clinical applications and copy number calling, we work with the history of copy number events represented as a CNA tree (Figure 1d). This is in analogy to the mutation trees of SCITE [39], where the tree nodes now encapsulate events corresponding to amplifications or deletions of the segments. The nodes in the tree can have arbitrary degree. We index the event nodes 1, …, *n* and additionally label them with the corresponding CNAs. All CNA events are stored in the vector *V*, such that *V*_*i*_ is the collection of CNAs of vertex *i*. For the example of Figure 1d, we have *V* = (+*S*_1_ + *S*_2_, +*S*_2_ + *S*_3_, −*S*_1_, − *S*_4_, +*S*_2_). The tree structure *T* we can store as a parent vector *T* = (0, 1, 2, 2, 1) where 0 represents the root.

If cell *j* is attached to event node *σ*_*j*_ we can read off the expected copy number state for each segment by tracing all events back to the root for a given tree structure and event vector. In the example of Figure 1a, the attachment vector is ***σ*** = (3, 3, 4, 1, 5). We denote the expected copy number state of cell *j* for a given attachment point *σ*_*j*_ as 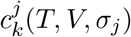. Then we have for the probability of the reads of cell *j* falling into segment *k*

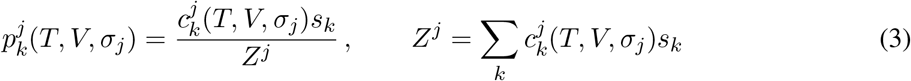

If a segment is entirely deleted so that the copy number state and probability is 0, then we would not expect to see any reads in that segment. However, due to mapping errors there may still be some residual reads in that segment. To account for this possibility, we instead set the minimum copy number state to 0 < *η* ≪ 1.

### Likelihood

To assess how well a CNA tree fits our read count data, we define a likelihood model as follows. The data matrix entry *D*_*jk*_ stores the number of corrected counts cell *j* has in segment *k*, with total reads for that cell of 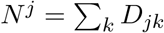. Given the probabilities of reads landing in each segment, we model the data with a Dirichlet-Multinomial distribution to account for overdispersion

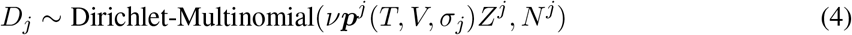

where *νZ*^*j*^ is the concentration parameter (inverse of the overdispersion). We scale by *Z*^*j*^ to aid the computational implementation since, using Equation (3), the likelihood contribution from cell *j* is therefore

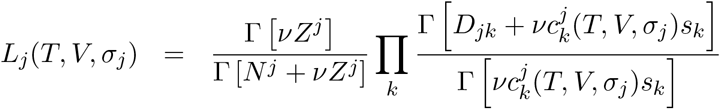

In the large *ν* limit, the model simplifies to the multinomial distribution

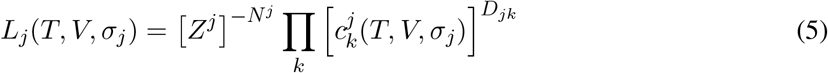

We can compute this likelihood for all cells and all possible attachment points efficiently in total time *O*(*mn*) where *m* is the number of cells and *n* the number of event nodes. This time complexity is achieved with a tree traversal. During the tree traversal, at each step only the segments in one event node change their state and we only update a limited number of terms in the product. All possible attachment points of a cell can therefore be computed in linear time.

For numerical accuracy, we compute the log-likelihood of all attachment points for each cell relative to the root. At the root, with a given constant ploidy *c*, we have the score per cell of

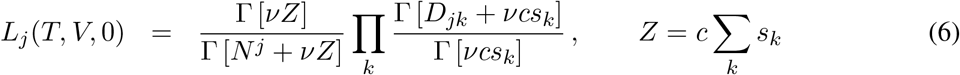

whereas for large *ν* the ploidy constant cancels and we simply have

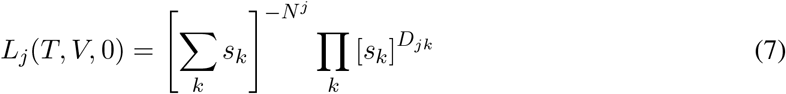

Since the root is common across all tree models, this contribution only varies if *ν* is changed and is only recomputed in that case. The root does not need to have constant ploidy; for example the sex chromosomes can be set accordingly.

### Posterior tree probability

With the likelihoods of each cell for each attachment point, we can directly compute the marginal likelihood when we average over all possible attachment points of each cell in the tree

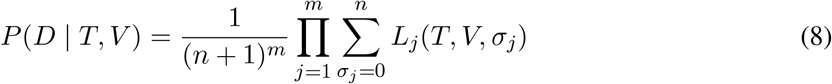

with a uniform prior on the attachment points. With a non-informative prior on the tree size, and then a uniform prior also on the trees, the posterior becomes

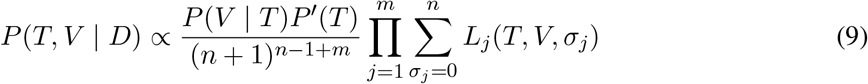

where *P*′(*T*) is a penalisation term needed to account for combinatorial effects possible for larger trees which we detail in Appendix B, and where we discuss the prior on the event vector in Appendix C.

Since we marginalise out the cell attachments, we focus on the CNA tree and build a scheme to find the tree and vector with the highest posterior score. Not all combinations of trees and event vectors are meaningful. After a copy number state of 0 is reached at any point in the tree, the affected segment cannot be regained in any descendant nodes. Also, a tree with a placement of events such that it recreates the exact same genotypes repeatedly in different parts of the tree is redundant, as a simpler model using a smaller tree would fit the data equally well. Biologically such convergent evolution may occur, but as it cannot be distinguished by the data itself, we make the assumption of the more parsimonious model. To exclude the possibilities above we assign any trees with forbidden events a posterior score of 0.

As an alternative to marginalisation, we can also place cells at their maximal attachment position and define the score

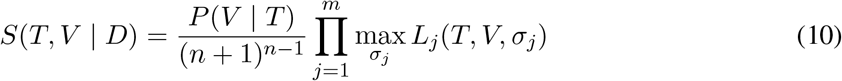

which removes the need for correcting for combinatorial effects via *P*′(*T*). To distinguish between these two alternatives, we denote inference using the marginalised score, Equation (9), with the term ‘sum’ and inference using the maximised score, Equation (10), with the term ‘max’ in the main text (for example in the simulation plots of Figure 5).

### MCMC

To sample trees proportionally to *P* (*T, V* | *D*) we build an MCMC scheme to move in the space of trees and event vectors. We detail the moves in Appendix D starting with the simple moves of *prune and reattach* and *label swap* employed in SCITE [39] which work for a fixed tree size and fixed set of events at the nodes. We additionally weight the move proposals to account for their computational costs. To change the event vector, we introduce an *add or remove events* move, while to change the tree size we developed the *add or remove node* and *condense or split nodes* moves to cover the full discrete space. To aid moving in such a complex space and finding simpler trees which generate the same set of possible genotypes we also introduce a *genotype-preserving prune and reattach* move. Since the set of genotypes is preserved, only the event vector prior needs to be recomputed making this move computationally very efficient, so we score and sample from the whole neighbourhood. For the overdispersion determined by the concentration parameter *ν* we run a random walk (in log space). For the complete MCMC scheme, at each iteration we first sample a move type with a fixed probability for the six different moves. Each move is equally likely to be chosen apart from the *genotype-preserving prune and reattach* which has a relative weight of 0.4.

### Maximal search

We utilise the MCMC scheme to search for the tree with the highest posterior score, keeping track of the maximally scoring tree encountered. From the best scoring tree discovered, we can compute the best attachment of each cell and hence their predicted copy number profiles.

To aid finding the best scoring tree and event vectors, we work stepwise. First we cluster the data (using PhenoGraph) and construct the average read counts per cluster. The tree likelihood computation only involves running over each cluster (weighted by the cluster size) rather than each cell, giving a large computational speed up. We initialise the states as the normalised read counts rounded to integers, and the tree by performing hierarchical clustering with Ward’s method and filling the events at the internal nodes with our extract common ancestor procedure. We run 10 different chains and check for robustness that at least half the chains return trees with similar high score (the log posterior is within 5 of the maximum). Non-robust runs are repeated starting from the highest scoring tree found so far. Once the tree on the clustered data has been determined, we run the scheme on the full data, starting from the cluster tree and with the overdispersion learned from the full data with the cluster tree. We compute the distances between the trees returned by half the chains that attain the best scores in the full data, and check for robustness that the average distance is less than 0.02. We stop the scheme on the full data once either of the robustness criteria are met.

### Root state

Instead of assuming a diploid state at the root, it is easy to start with any other state, for example tetraploid corresponding to a whole genome duplication. Although the average ploidy is not well defined for read depth data, because CNAs are integer, the differences induced can help determine the underlying states. Under the presence of further CNAs, the model comparison of the tree with a diploid root against that with a tetraploid root potentially allows identification of genome doubling relevant for tumour progression and prognosis [50]. We identify such a doubling in the 10x Genomics breast cancer dataset.

### Simulation settings

For the simulation we consider a genome consisting of 10,000 bins and different numbers of reads per cell with an average of 2, 4 or 8 reads per bin. This is a relatively low read depth per bin, chosen to make the inference more challenging and comparable to the data from the 10x Genomics protocol with their default bin size of 20kb.

For trees with size *n* = 20, we partitioned the bins into 40 or 80 segments. At each node in the tree we sampled the change in copy number by first sampling the number of segments involved with a Poisson distribution with parameter *λ* = 0.1 and added 1, while for each segment we sampled the number of copies with a Poisson distribution with parameter *λ* = 0.2 and again added 1 and sampled the sign uniformly. Trees that violate biological constraints (for example having negative copy numbers states) are rejected and resampled. For each setting, we sampled 40 trees.

For each tree, we sample 400 cells, in line with the typical number of cells targeted with the 10x Genomics protocol for a single sequencing run, and sample their attachment point in the tree uniformly. This provides the true copy number profile of each cell, from which the reads per bin were sampled with a Dirichlet-multinomial with concentration *ν* = 4.

For inferring the copy number profiles with our MCMC scheme, we first detected the breakpoints automatically (Appendix A with *ν* = 1) to generate the segments. For our MCMC scheme we set *η* = 10^−4^ and ran chains of length 4,000*n*, repeatedly until robust results were obtained, to find first the trees on the clustered data and then on the full data. We find the best trees both for the posterior score from the summation of Equation (9) and for the maximisation of Equation (10).

For comparison we include HMMcopy [51, 19] which runs a hidden Markov model on each cell to call the breakpoints and the copy number states which are inferred per bin for each cell individually. For single-cell low-read count data, however, default parameters of HMMcopy should be adjusted and we followed the recommendations of [26] (log-transformed counts, *e* = 1− 10^−15^, *ν* = 4, strength of 10^7^). We also include Ginkgo [20], which offers single-cell copy number calling, adapted to deal with cell-by-bin read count matrices and with the baseline, maximum and minimum ploidy set to 2, 6 and 0 respectively, and SCOPE [23], which we initialise by setting the top 10% cells with lowest Gini coefficient as diploid, and allow at most 20 copy number clones.

We further consider CONET [35], a tree-based method where the main difference between SCICoNE and CONET is that CONET does not explicitly model integer copy number changes in their tree nodes, but rather models breakpoint presences or absences as the events in the tree. Copy numbers are called in a post-processing step of mapping to the nearest integer, and while the breakpoints share an evolutionary history, the copy number states are not assumed to, unlike in SCICoNE. The input data for CONET is correspondingly not a cells by regions matrix, but rather the difference in normalised read counts between subsequent bins. For the simulations we followed the recommendations in [35] and set *k*_0_ = 1, *k*_1_ = 0.01, *s*_1_ = 100000, *s*_2_ = 100000 and *n*_*CN*_ = 2, and set the number of parallel tempering iterations to 200,000. In case the method took more than 24 hours to run, we reran with their default setting of 100,000 iterations. CONET also requires candidate breakpoints as input for which we used the scheme provided by their software which relies on CBS [52] and MergeLevels [53] for genome segmentation.

We additionally include a diploid profile and two clustering methods. For the clustering we use both PhenoGraph [44] and hierarchical clustering (with the number of clusters chosen according to the Calinski-Harabasz index) and assigned all cells in each cluster the average read counts of that cluster. For each cluster and its associated cells, we then set the median counts per cell to 2 (by doubling and dividing by the median) assuming that diploid is the most common state and rounded the copy number states to the nearest integer. In the comparison we compute Δ as the root mean squared difference over all cells and bins, between the true copy number profiles and the inferred values.

For the comparison, we used the distance Δ between the inferred and simulated copy number states, rather than a distance between the structures of the inferred and simulated trees since none of the alternative methods generates a CNA tree. The distance Δ works on the set of predicted genotypes, so that errors in the tree structure which generate incorrect genotypes for cells will be picked up by this distance measure.

As a further measure we adapt the path difference metric [54], previously used for clone and mutation trees [55, 39], to CNA trees. Specifically we define a distance between a pair of cells on tree *T* along the shortest path between the cells through their most recent common ancestor. Along the path we count the total number of events encountered, with each event weighted by the number of bins it affects. Equivalently for each bin we count how many times it changes state, and by how much, along the path and sum over all bins:

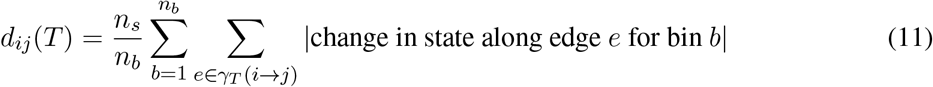

where *n*_*b*_ is the number of bins, *n*_*s*_ the number of segments and *γ*_*T*_ (*i* → *j*) the shortest path between cells *i* and *j* on tree *T*. We normalise the distance to count in units of ‘typical’ events by scaling by the expected size of an event 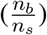. To compare an inferred tree *T*′ to the true generating tree *T* we compute the root mean square difference between the (triangular) distance matrices for the two trees

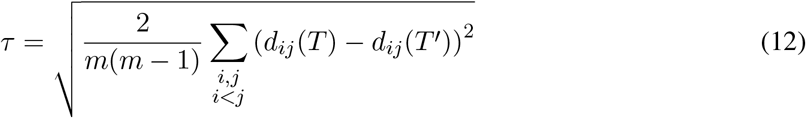

When log-transforming *τ* for plots, to avoid numerical issues we add an offset of 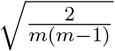 corresponding to a difference of a single event between one pair of cells.

## Code availability

SCICoNE has been implemented in C++ and is freely available under a GNU General Public License v3.0 license at https://github.com/cbg-ethz/SCICoNE. The code repository also provides a Python package to embed SCICoNE with data pre-processing steps and downstream analyses of the results, and includes an interface for data generated with CellRanger. We additionally provide Snakemake [56] workflows to reproduce all our results.

## Data availability

The xenograft sequencing datasets utilized in this study were generated by [16, 42] and are available at the European Genome-phenome Archive (http://www.ebi.ac.uk/ega/) under accession number EGAS00001002170 and EGAD00001004552. The dataset from 10x Genomics can be downloaded from https://support.10xgenomics.com/single-cell-dna/datasets as ‘Breast Tissue nuclei section E 2000 cells’.

## Acknowledgements

J.K. was supported by ERC Synergy Grant 609883 (http://erc.europa.eu/), P.F.F. by SNSF Grant 310030 179518 (http://www.snf.ch) and K.J. was supported by SystemsX.ch RTD Grant 2013/150 (http://www.systemsx.ch/).

---

**Figure A1:**
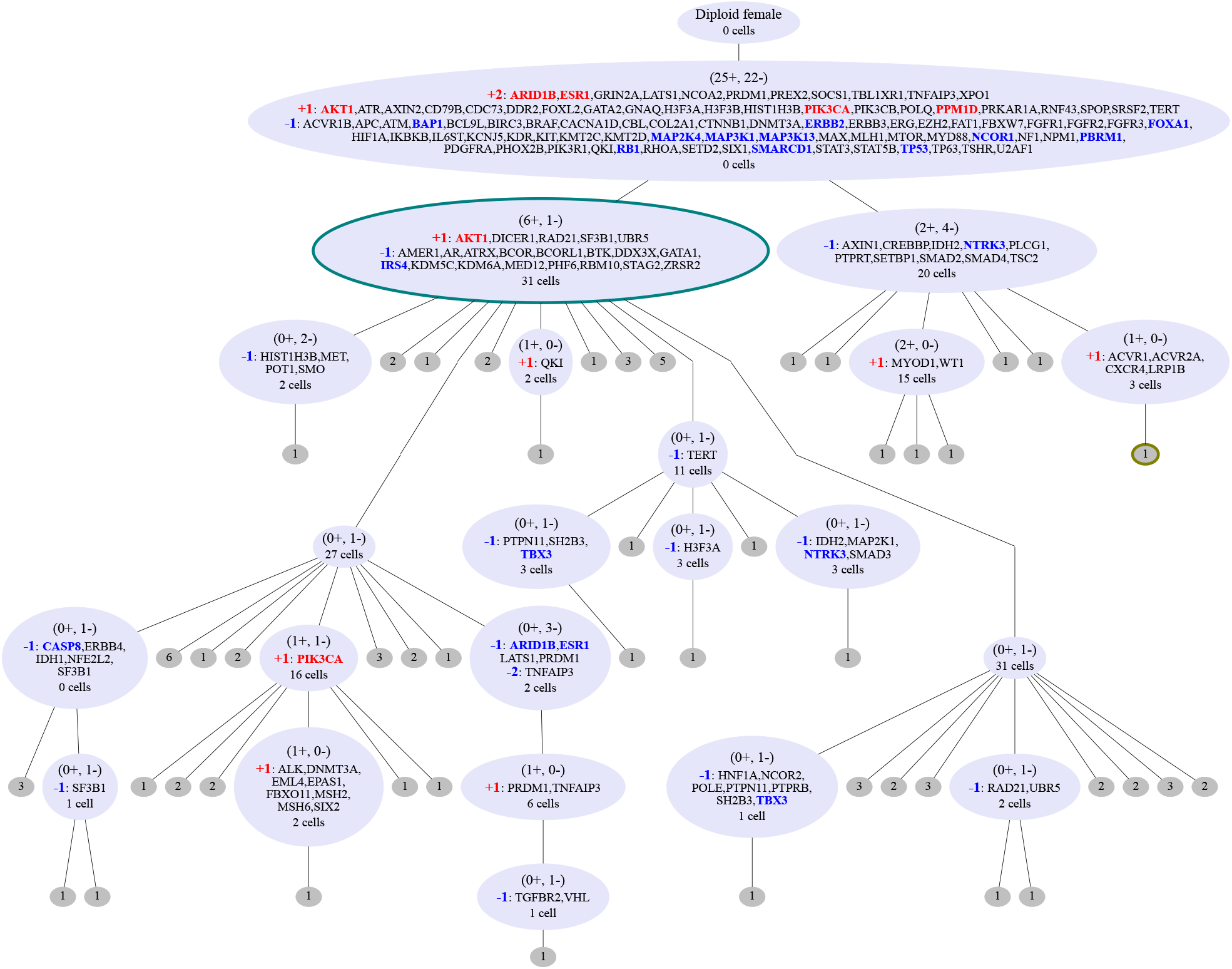
Inferred tree for 260 cells from a breast xenograft [16]. An expanded version of Figure 2 showing all the genes from the COSMIC Cancer Gene Census [41], indicating where and by how much they are amplified or deleted. Highlighted are the subset of 33 genes associated with breast cancer displayed in Figure 2.

**Figure A2:**
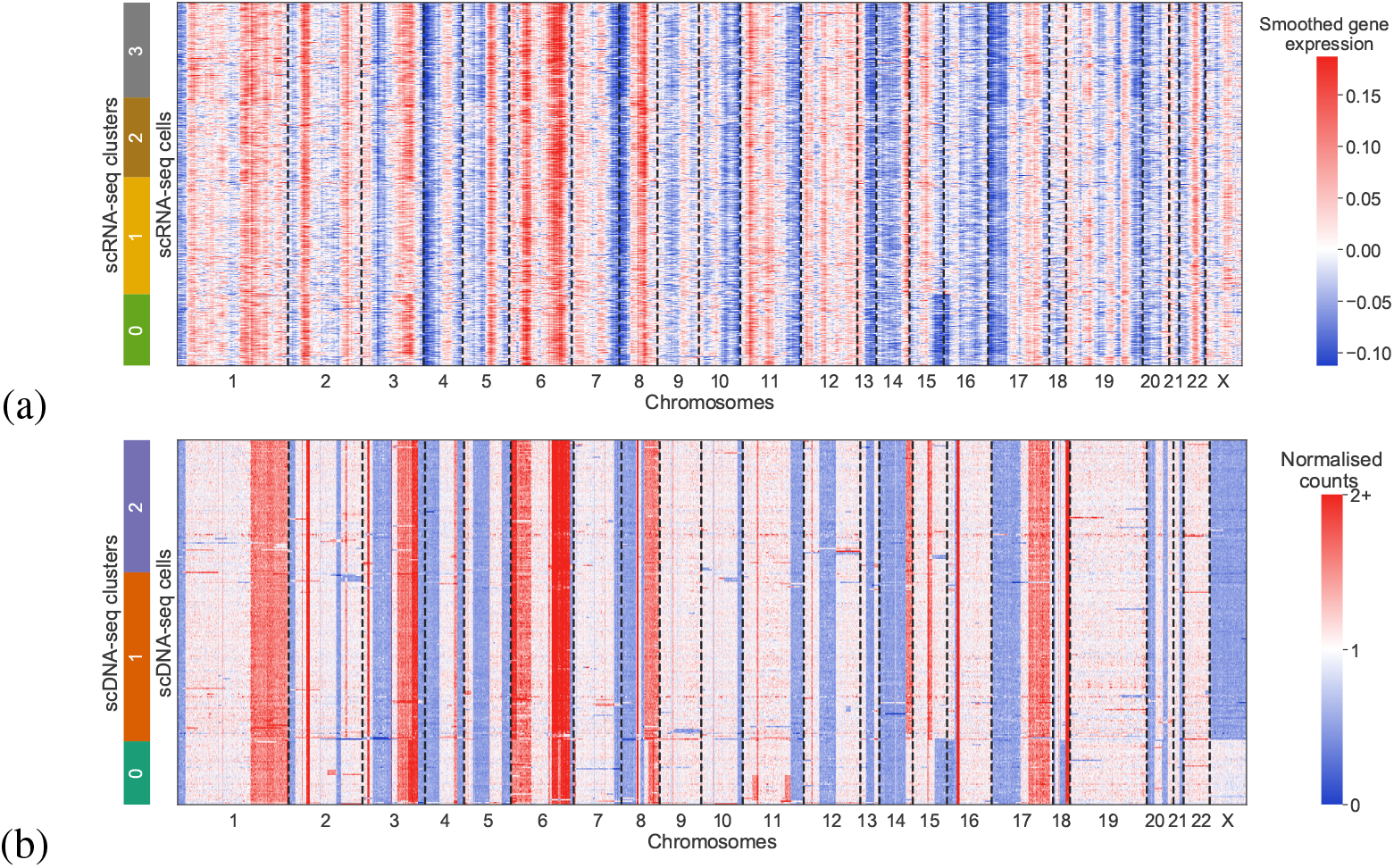
RNA and DNA levels for breast xenograft cells. (a) The smoothed expression profiles of 1152 cells from single-cell RNA sequencing [42]. (b) The normalised counts per gene for the expressed genes for 260 cells from single-cell DNA sequencing [16] are displayed for comparison.

**Figure A3:**
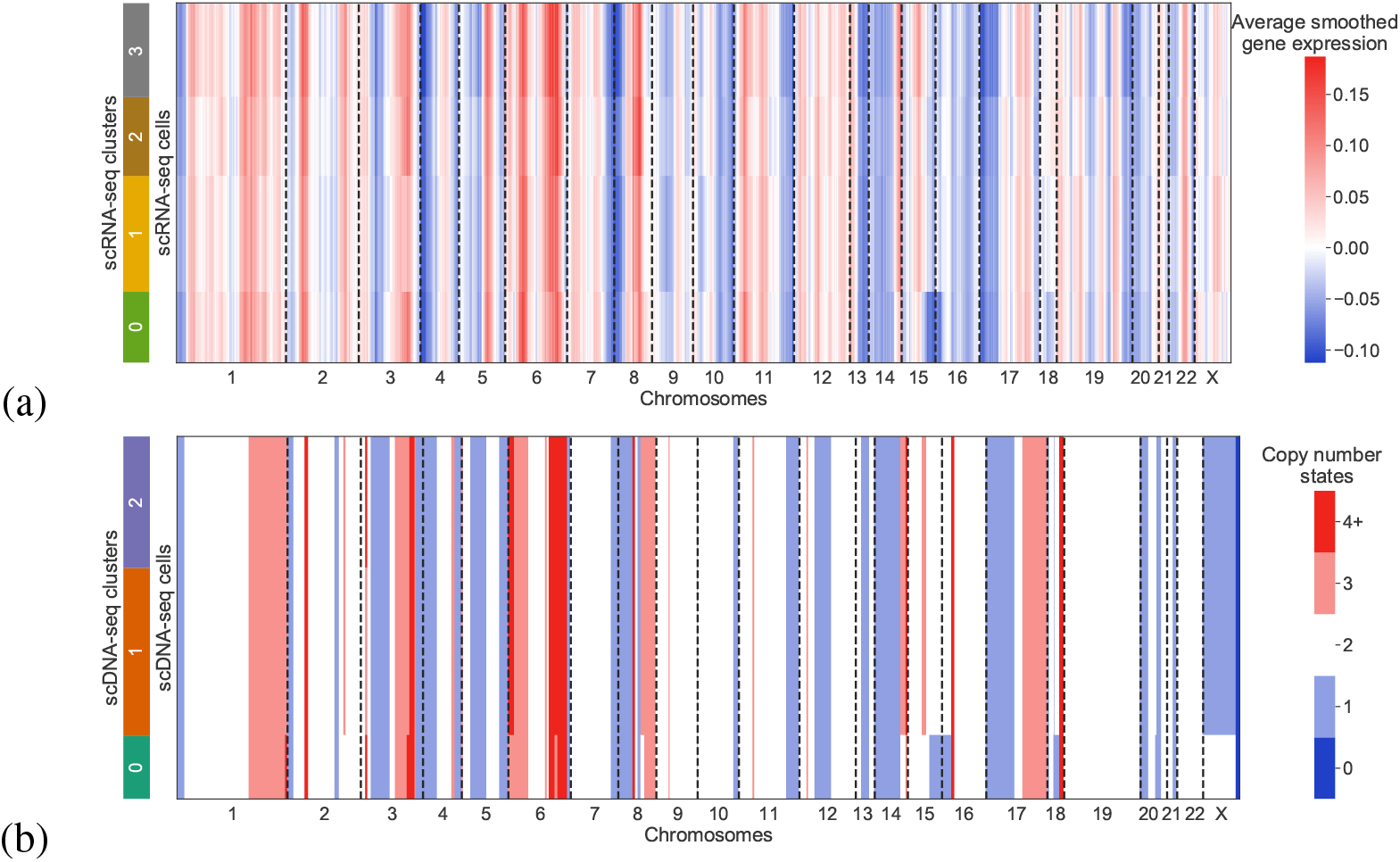
RNA level and DNA copy number for breast xenograft clusters. (a) The smoothed expression profiles of the cells in Figure A2a are clustered and the average profile of the cluster displayed. (b) The inferred copy number profiles of clones built from clustering the single-cell DNA sequencing data and learning a tree using SCICoNE (displayed at the gene level to match the normalised counts in Figure A2b). The inferred copy number profiles of the full data are in Figure 3.

**Figure A4:**
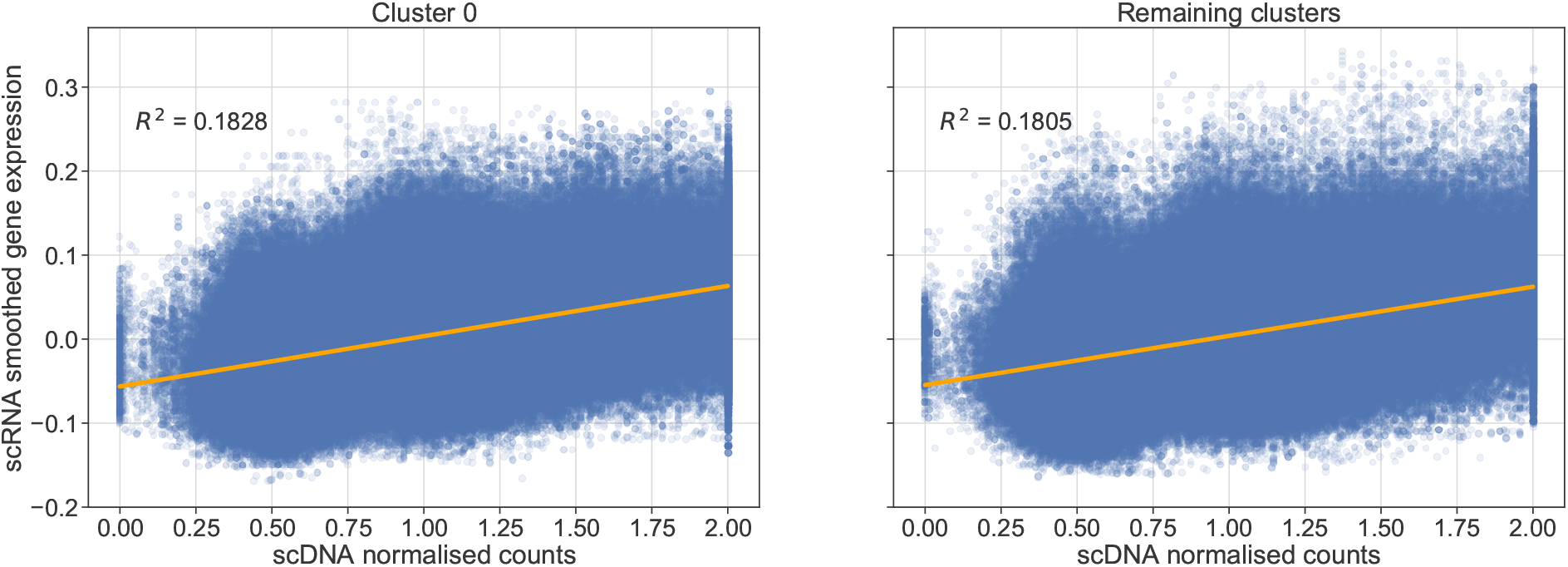
Correlation between RNA and DNA profiles. For the smoothed RNA expression and normalised DNA count profiles (Figure A2), we look at the correlation between the two modalities. Since different cells underwent each sequencing, we compare an RNA cell to a DNA cell both from cluster 0 (left panel) or both from other clusters (right panel) and plot the RNA expression level and DNA count depth for each gene. This is repeated for 100 random pairs of cells to obtain each scatter plot and correlation line.

## Appendices

### A. Breakpoint detection

For a given bin position *ρ*, we wish to test if there is a change in read counts in the next bin, across cells. In particular we fix a window size *ω*, consider the bins from *ρ* − *ω* + 1 to *ρ* + *ω* and look at the evidence for a breakpoint, and hence a copy number change, after bin *ρ*.

#### A.1 Evidence per cell

For a given cell *j* we model the read counts in bin *i*, 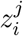 with a negative Binomial to account for overdispersion. We parametrise in terms of the mean *λ* and overdispersion parameter *ν* with a mass function of

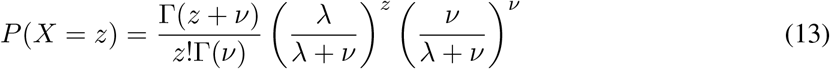

If there is no copy number change after bin *ρ* we would have equal expected counts across all bins in the window, so that the log-likelihood of the observed counts is

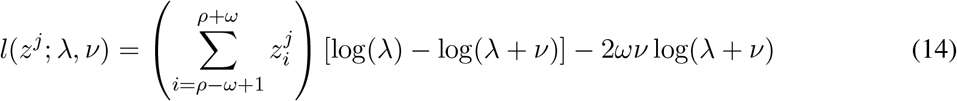

where we ignore constant terms that do not depend on *λ*. The maximum likelihood, for fixed *ν* occurs at

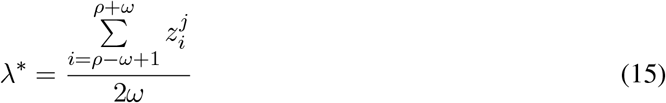

To allow for noise giving unbalanced counts on either side of *ρ* even without a breakpoint, we fit a linear model for the expected counts

**Figure A5:**
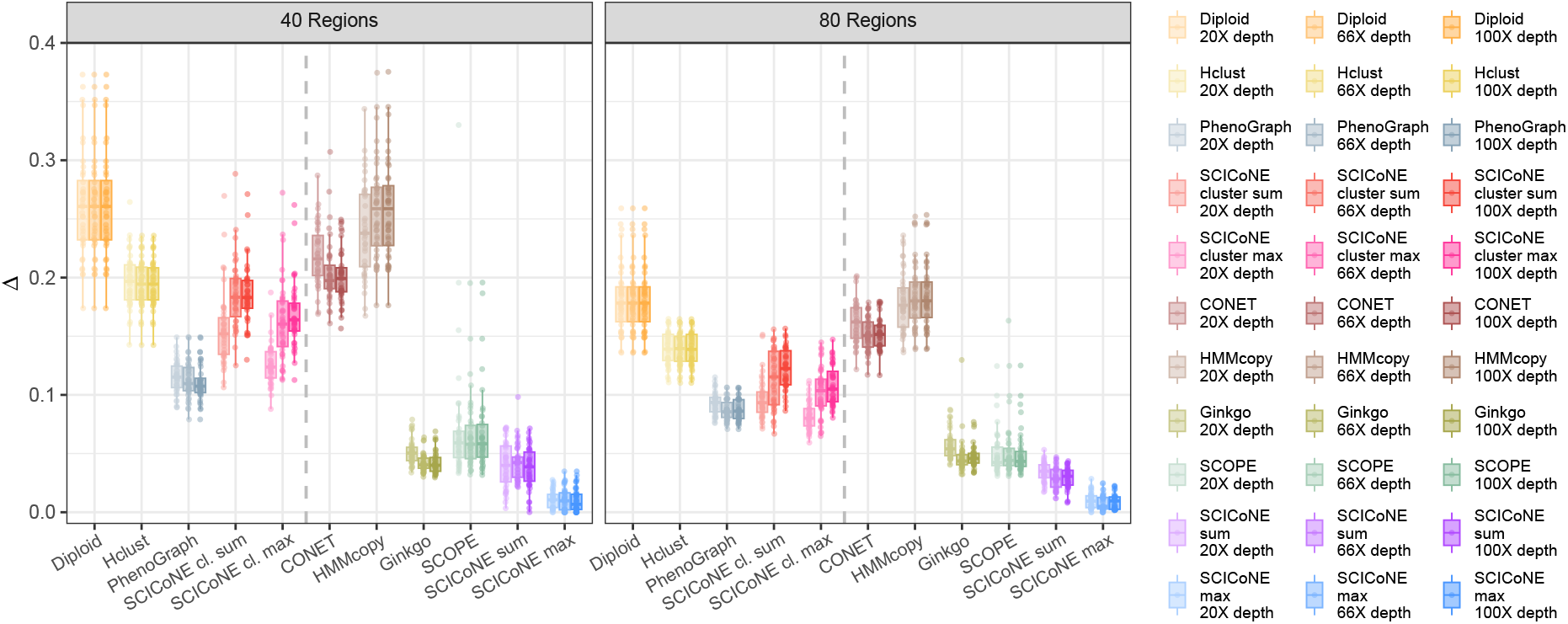
Comparison of copy number calling for simulated data. The simulation setting is as in Figure 5 but with higher coverage and without overdispersion.

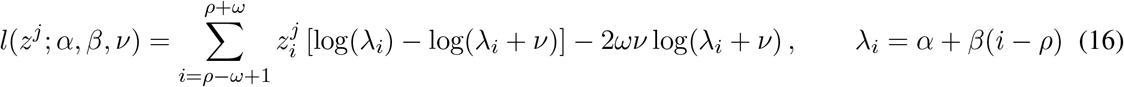

which we maximise over *α* and *β*:

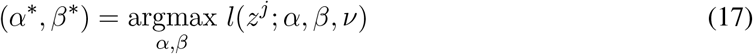

To avoid fitting real copy number changes with the linear model, we bound the slope so the relative change across *ρ* is less than a quarter by restricting 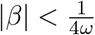.

If there is a breakpoint, we would expect to have different average counts on each side which we model with two mean parameters leading to a log-likelihood of

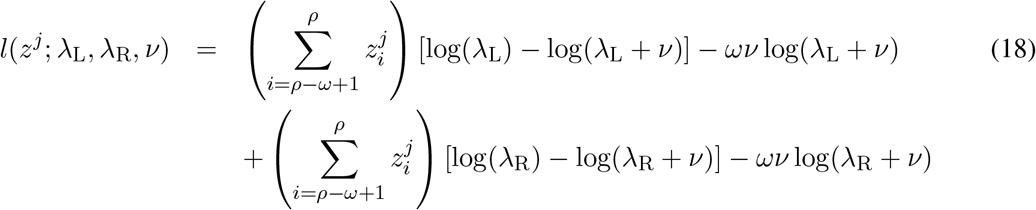

This is maximised by the average counts of each side of the potential breakpoint: For robustness and to uncover changes in count level that spread over many bins, we use the robust mean

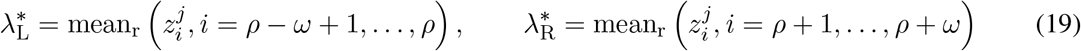

We further enforce a minimum difference between the *λ* parameters on each side of *ρ* so that the relative change in copy number is more than a quarter:

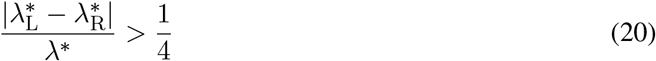

If the inequality is not satisfied by 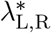, we rescale their differences from *λ*^*^.

Finally we compute the difference in maximum log-likelihoods of the two models. We compute this difference per cell for each breakpoint

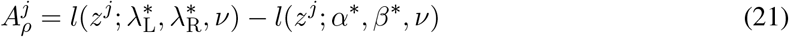

Especially at low read-depths, the likelihood ratios over bins 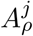 for each cell may exhibit noise from the underlying count data. However we would expect breakpoints with a copy number change to provide a signal across the bins with a width of the window size *ω*. To amplify these signals and filter out noise, we perform a low-pass Gaussian filter (with half-gain cut-off frequency corresponding to twice the width *ω*) with a Fourier transform and replace the 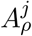 signal per cell by the smoothed version. Since this filter may help identifying true breakpoints with high-noise, but by adding width to the signal may add uncertainly to their detected location, by default we only employ the filtering for read depths of 10 and below.

#### A.2 Combining cells

Next we compute the logarithm of the combined evidence that the break-point occurred in any *k* of the *m* cells with the average

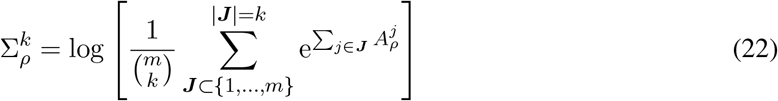

This can be computed efficiently using dynamic programming (Algorithm 1). We further combine the evidence for the breakpoint in *k* of the *m* cells with the prior of an event affecting *k* cells in a random binary cell lineage tree [27]:

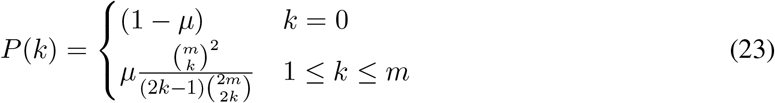

where *µ* is the prior probability of an event occurring. The posterior probability of the event occurring in *k* cells is then proportional to 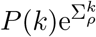. To rank the breakpoints, we consider the posterior probability of the breakpoint occurring in *k*^*^ or more cells

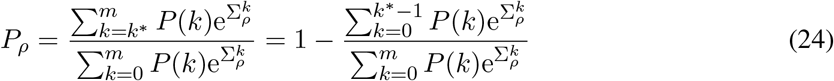

and compute

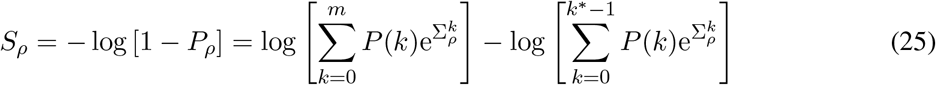

#### A.3 Peak detection

Plotting *S*_*ρ*_ across the genome we find peaks, with a width of *ω* when *ρ* lines up with a breakpoint in a number of cells (Figure A6). Since the peaks possess very different heights, we perform a further log transform, 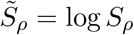. Since the local median value of 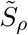 may depend on the underlying copy number state, we find the breakpoints iteratively. First we divide 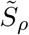 by its median value and find the highest value. If it is above the threshold, we split 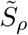 into two parts, exclude a window of size *ω* from either side and divide the remaining parts by their median. We repeatedly find the maximum value, split the vector excluding a window around the maximum, and divide the regions between the current breakpoints by their median until the maximum in those regions is below the threshold. The threshold has a default value of 3 times the distance between the median and third quartile of the bins not around the current set of breakpoints. We have a default window size of 20.

##### Algorithm 1

Obtain the sum of breakpoint evidence over all cell subsets

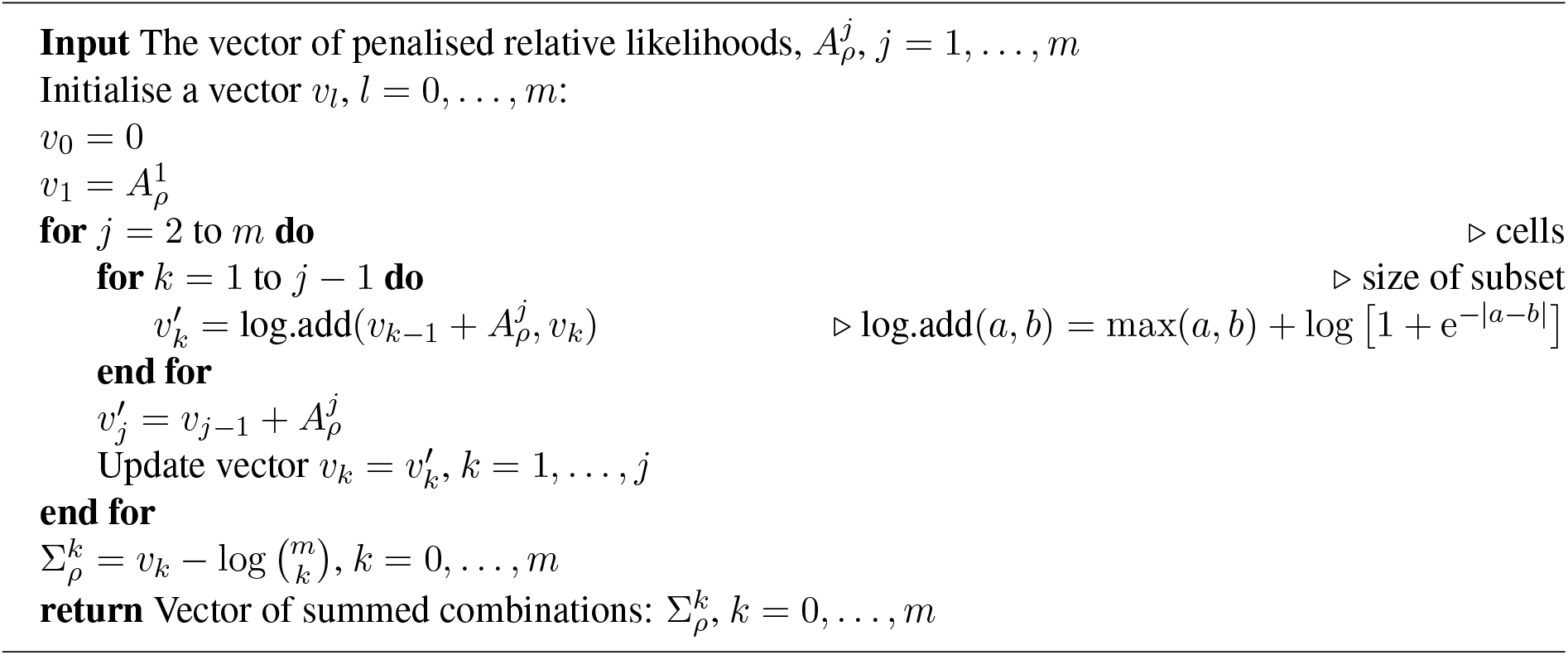

**Figure A6:**
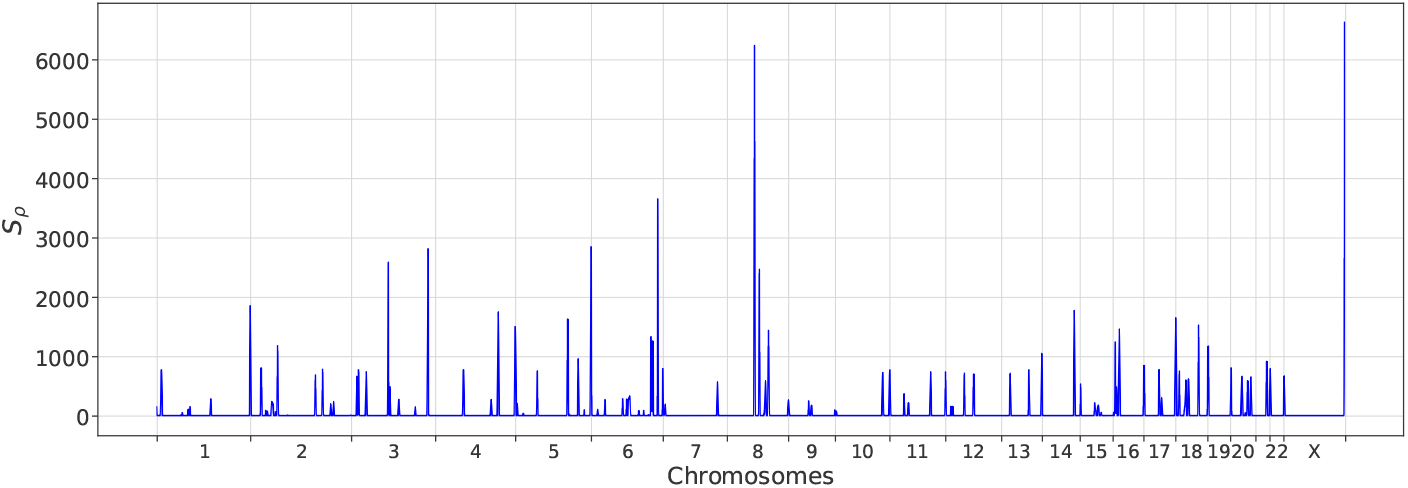
Breakpoint detection. For each bin we combine the evidence for a breakpoint across the 260 cells from a breast xenograft [16], to arrive at *S*_*ρ*_. Peaks corresponds to putative breakpoints.

### B Tree penalisation for combinatorial effects

For any tree, assume that the largest cluster of cells of size *m*_*c*_ attach best to the same node *k* and poorly elsewhere. The likelihood contribution of these cells includes the term

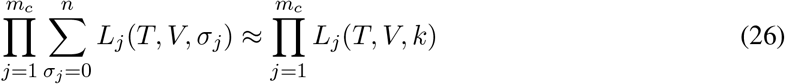

where in the sum we assume that the other attachment points are negligible for the sum. Now imagine we add another node *k*′ to the tree with a very similar genotype to the best attachment point of the cells, and hence very similar likelihoods for the cells. The likelihood term now becomes

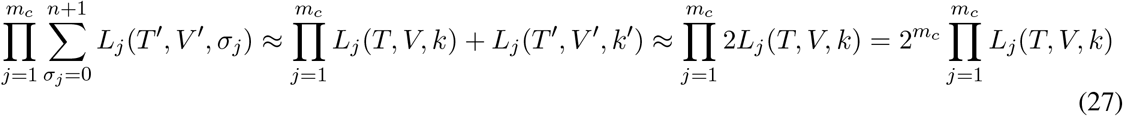

so that it can be increased by a factor of up to 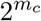. The likelihood contribution can be made arbitrarily large by adding further dummy nodes with similar genotypes. To counteract this effect, we can penalise trees with the factor

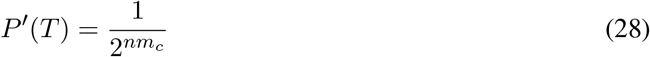

so that if we add an additional node and increase *n* by 1 we obtain an additional factor of 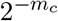.

### C Prior on the event vector

For the prior on the event vector *V*, we simply consider that each amplification or deletion may be selected among the *K* segments with a choice of sign, leading to a factor of 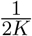. At each node, we consider the amplification of contiguous segments as a single amplification event, and likewise for deletions. To compute this number, let us list as ***v***^*i*^ a vector of the number of copy number events for each segment occurring at node *i* in the tree. For the example in Figure 1c with event vector *V* = (+*S*_1_ + *S*_2_, +*S*_2_ + *S*_3_, −*S*_1_, −*S*_4_, +*S*_2_) we would have ***v***^2^ = (0, +1, +1, 0, 0) over the *K* = 5 segments. To count the number of contiguous amplifications and deletions we can simply count when the copy number profile increases as

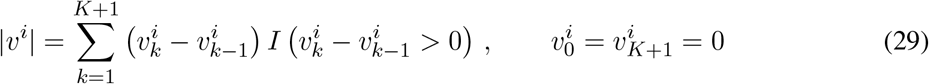

where *I* is the indicator function. The prior probability for each node is then 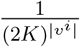.

For the full event vector prior, we may freely permute the ordering of the event vector, along with the corresponding numbering of the event nodes in the tree. We therefore divide by the overcounting factor of *n*! to arrive at

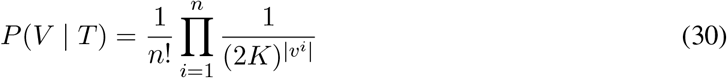

Note that although the overcounting factor assumes that the elements of the event vector are distinct, we keep the same value in all cases for simplicity.

As a further penalisation, whenever an event has the opposite sign to the cumulative copy number state of the parent node, we subtract *α* from the log-likelihood, for which we have the default value *α* = 10.

### D MCMC moves

#### Prune and reattach

The most basic move to change trees with a fixed event vector is *prune and reattach*. From the current tree *T*, we propose a new tree *T*′ by uniformly sampling a node (except the root), detaching it and its descendant subtree and then uniformly sampling a new parent from the remainder of the tree including the root. An example of this proposal move is depicted in Figure A7. After the new tree has been proposed, we simply accept the move with probability

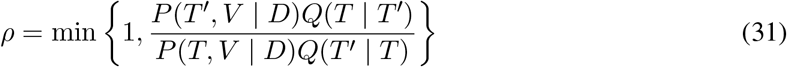

where *Q*(*T*′ | *T*) is the probability of proposing tree *T*′ when currently at tree *T* in the chain. Since the move is symmetric, *Q*(*T*′ | *T*) = *Q*(*T* | *T*′), the acceptance ratio simplifies to

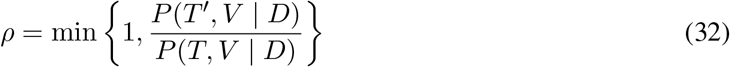

Since only the detached subtree needs to be rescored, it is cheaper to move smaller subtrees and more of such moves can be processed for the same computational cost. To try and speed up convergence of the MCMC scheme, we can preferentially sample nodes to move which have fewer descendants. We denote by *d*_*i*_(*T*) the size of the subtree of *T* starting with node *i* (the number of descendants plus 1). Instead of uniformly sampling nodes, we can sample proportionally to the *d*_*i*_(*T*)^−1^ to balance the computational cost of different moves.

**Figure A7:**
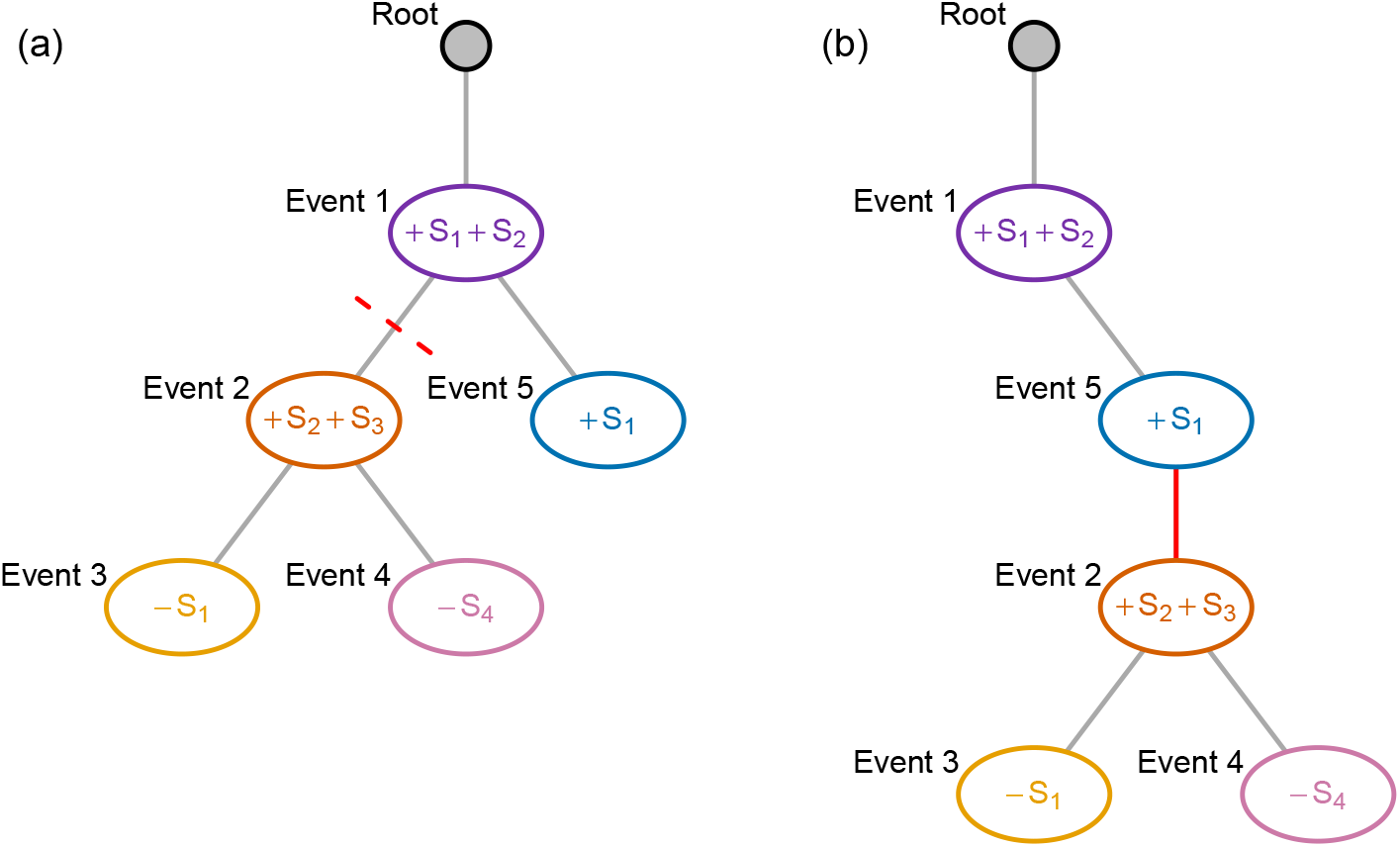
Prune and reattach. From the event tree in (a), we propose the new tree in (b) with the *prune and reattach* move by sampling a node (event node 2), detaching it from the tree and sampling a new parent (node 5).

If we define

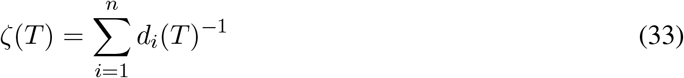

as a normalising constant then the transition probabilities in moving from tree *T* to *T*′ where node *i* is selected to detach are then

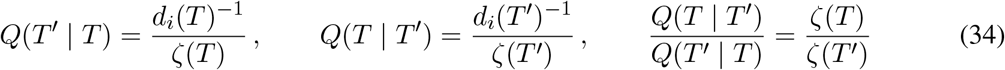

and the acceptance probability becomes

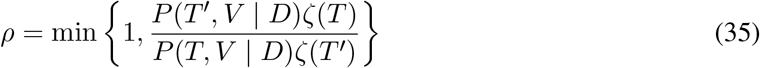

#### Swap node labels

A further move is to select two nodes uniformly at random and swap over the events associated with each node. All nodes below either affected node need to be rescored.

This move is symmetric and hence accepted with probability

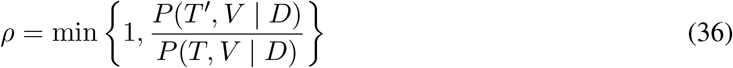

However, again we create a weighted version. We define *e*_*ij*_(*T*) to be the number of nodes affected by the swap. This is the number of descendants of node *i* plus the number of descendants of node *j* plus 2, if *i* and *j* are in different lineages, or the number of descendants of the ancestor minus the number of descendants of the lower node if they are in the same lineage. Then we can sample the pair (*i, j*) proportionally to *e*_*ij*_(*T*)^−1^. Since the tree structure does not change however, this move is also symmetric.

**Figure A8:**
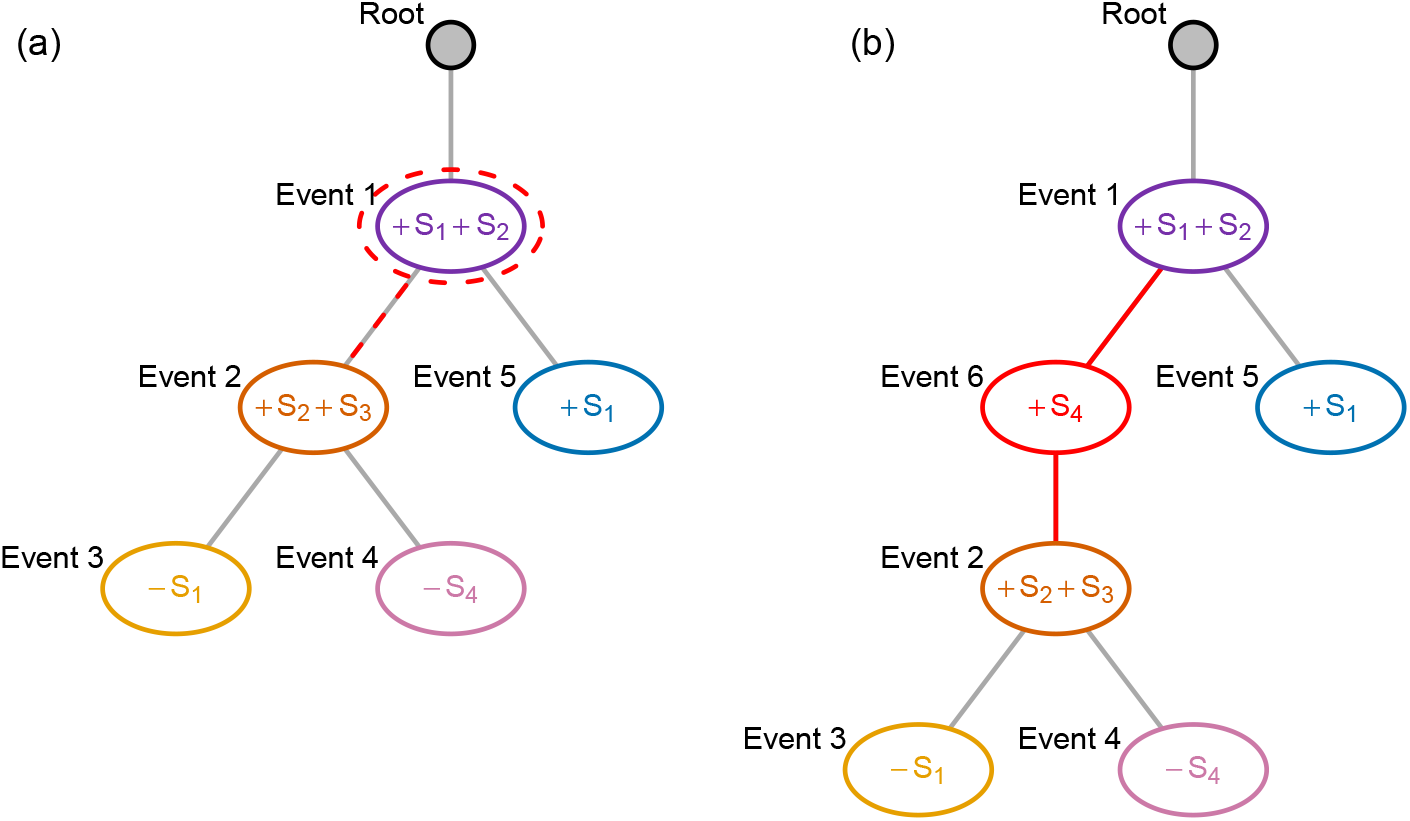
Add or remove node. From the tree in (a) [which is the tree in Figure A7a] we sample to add a new node and select event 1 as the new parent. From the two children of event node 1, we select event node 2 to become a child of the newly inserted node. With the node inserted, we sample the events in the new node as an amplification of segment 4 to arrive at the tree in (b).

#### Add or remove events

This move changes the event vector, but keeps the tree fixed. We first select a node at random (uniformly). Then we sample the number of segments to be affected from a Poisson(*λ*_*R*_) + 1 and sample this number of (distinct) segments uniformly. Then for each segment we sample the number of additional copies from a Poisson(*λ*_*C*_) + 1 and a sign uniformly. If the change leads to all events at the node being completely cancelled (say at node 3 in Figure A7a which only has the event −*S*_1_ we sample adding +*S*_1_), then we simply reject the move.

This move is symmetric (since the sign accounts for adding and deleting) so the acceptance ratio is

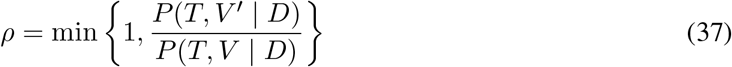

Setting *λ*_*C*_ = 0 and *λ*_*R*_ = 0 makes this move select only a single segment, and only allows its number of copies to change by 1.

In the weighted version, we sample nodes proportionally to *d*_*i*_(*T*)^−1^ as for the weighted prune and reattach, but since the tree structure does not change, the move is still symmetric.

#### Add or remove node

This move changes both the tree structure and the event vector, as in the example of Figure A8. To define the move, we first consider the possible ways of adding a node. We may place the new node below any of the (*n* + 1) in the current tree including the root. However, when we place the new node below a node withδchildren, any of theδmay become children of the new node instead of remaining siblings. There are therefore

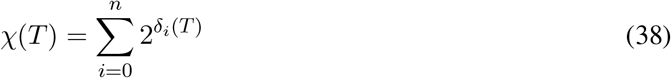

possible placements of the new node in the tree *T* where *δ*_*i*_ is the number of children of node *i*. For the example in Figure A8a thenδ= (1, 2, 2, 0, 0, 0) and hence *χ* = 13.

For the new node, we sample the number of segments to be affected with a Poisson(*λ*_*R*_) + 1. For each of the *r* segments we select from the *K* segments in total with 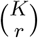 ways. If *r > K* we just reject the move for simplicity. Then for each of the selected segments we sample a sign uniformly and a number of copies *c* from Poisson(*λ*_*C*_) + 1. The transition probability, given that we selected to add a node, to that exact tree with that exact label is then

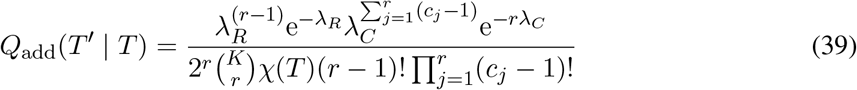

Since there are many more ways to add a labelled node, than to delete one, we weight the deletion by the corresponding terms in adding the node back. For example, for each node *i* of the *n* nodes to delete, we count the number of segments *r*_*i*_ and the number of copies *c*_*j*_, *j* = 1, …, *r*_*i*_ and weight that node by the factor

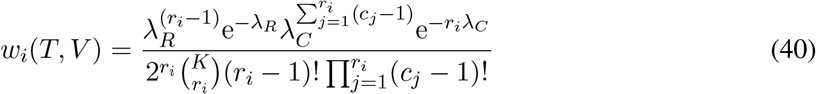

with the sum of these deletion weights, 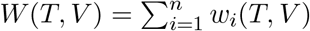.

For the move, we first sample, with equal probability, whether to delete or add. If delete is chosen then we sample a node from the *n* available proportionally to the weights *w*_*i*_(*T, V*), with probability 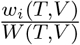. The sampled node is removed and all of its children are attached to the nodes previous parent. If add is chosen, we sample a node from the (*n* + 1) available including the root proportionally to 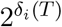. The new node is inserted as a child of the sampled node. The previous children of the sampled node are each independently randomly assigned to become children of the new inserted node with probability 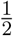, otherwise they remain siblings of the new inserted node. Finally the label of the inserted node is sampled with the Poisson distributions for the number of segments affected, their numbers of copies and uniformly chosen signs for each.

Since the weights for deletion include all the terms corresponding to sampling, for detailed balance we accept the move with a probability of

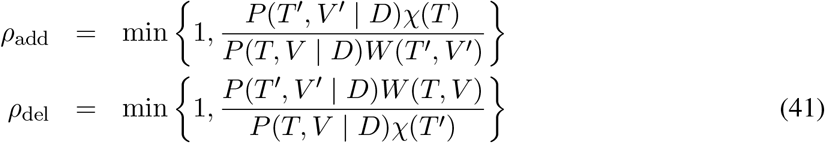

depending on whether add or delete was selected.

For weighting the moves also according to their computational cost, we need to further weight each term in the proposal neighbourhood by the cost of updating the score. For each node *i* to be deleted there are a remaining (*d*_*i*_(*T*) − 1) nodes to rescore so we define a new weight (without the minus 1)

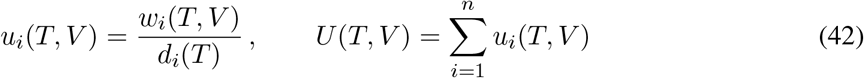

For adding nodes, say the new parent has *δ* children, we would need to compute the weights for all 2^*δ*^ choices. For simplicity we weight all those choices by assuming that on average only half of its descendants are affected along with the new node. The weights for adding are

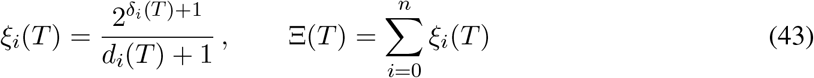

and the acceptance probability becomes

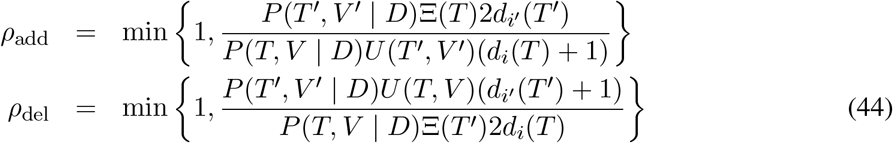

**Figure A9:**
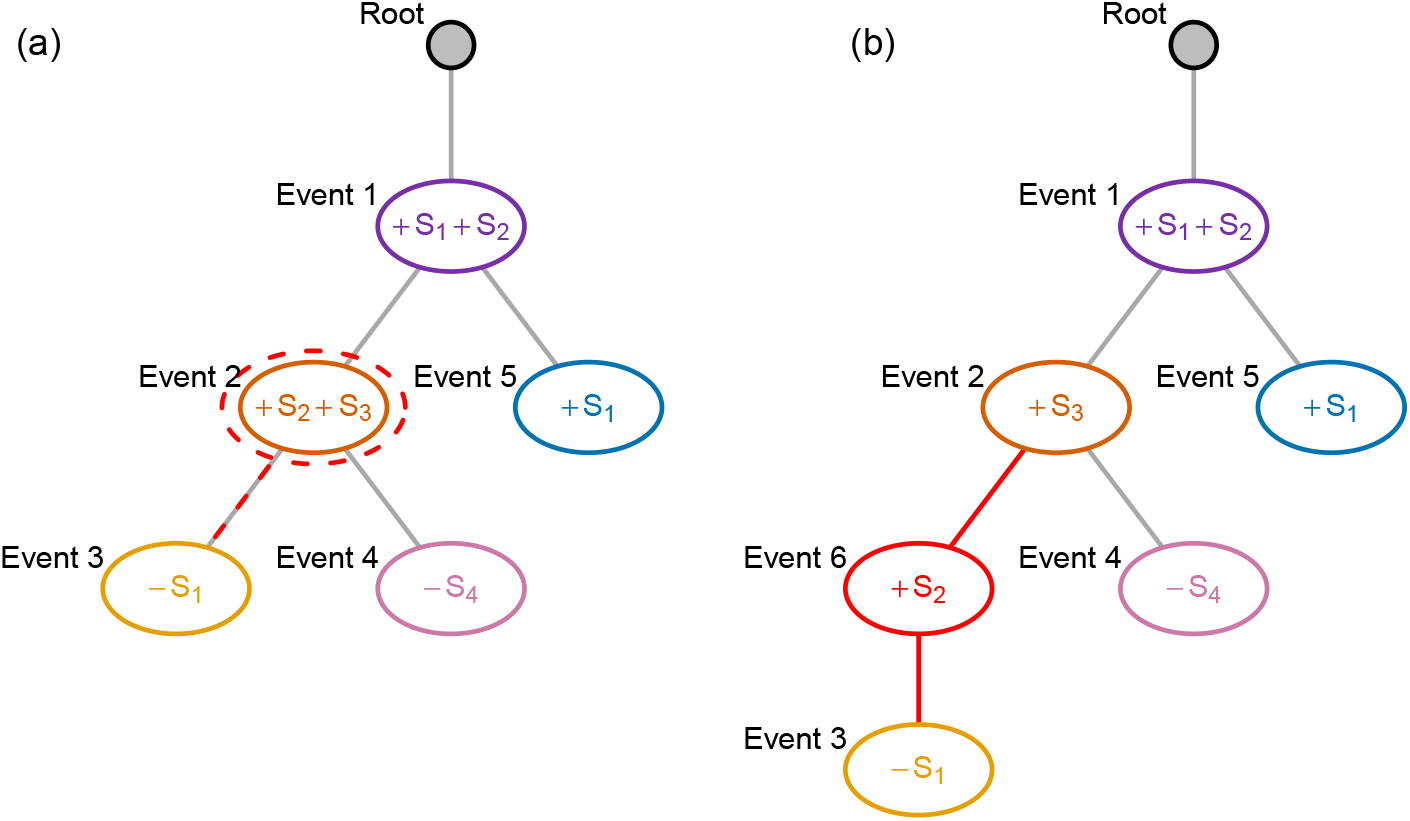
Condense or split nodes. From the tree in (a) [which is again the tree in Figure A7a] we sample event node 2 to split. From its two children, we select event node 3 to become a child of the newly split off node which becomes event node 6. With the node placed, we sample how to split the events which were previously at event node 2 across that node and its new child to arrive at the tree in (b).

#### Condense or split nodes

Finally we allow nodes to be split into two, or to combine a parent and child into one, as in the example of Figure A9.

We first consider the move where we split a single node into a parent-child. If we select the node *i* we would look at the vector ***v***^*i*^ of the copy number events at that node. Let’s say we have the vector ***v*** = (0, − 1, 2, 0, 0) which we need to split into two vectors. We sample the splits with a Poisson distribution with parameter *λ*_*S*_ in the following way:

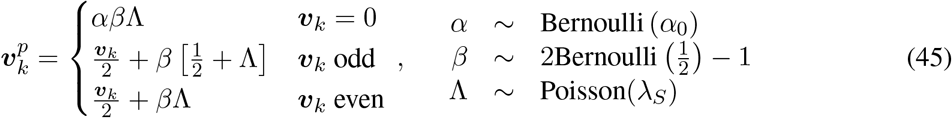

where for positions which are 0 in ***v*** we allow them to be split into cancelling events with a low probability *α*_0_. This sampling provides the proposed event vector of the parent, while that of the child is fixed by ***v***^*p*^ + ***v***^*c*^ = ***v***. If either vector ends up being completely 0, we reject the move.

For each of the segments we compute the absolute difference *c* between the number of copies in the new parent vector and the new child vector. The probability of picking each difference is

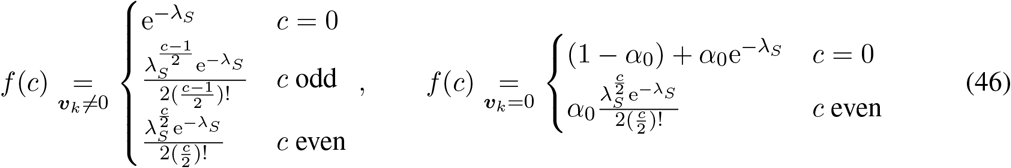

For the split, we do not consider the root so there are

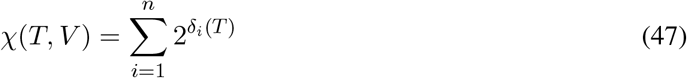

ways of arranging splits. The transition probability to that exact tree and exact event vector is

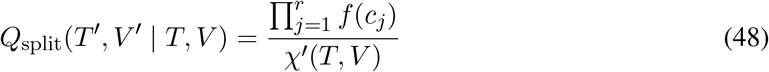

To compensate for this we weight the reverse move of combining a node with its parent by

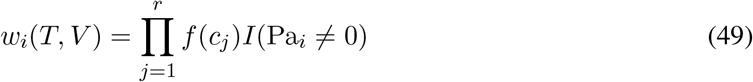

where the *c*_*j*_ are the absolute difference in copy number change in the non-zero segments in the node and its parent. Nodes attached to the root get a weight of 0, which we denote with the indicator function *I* on the parent Pa_*i*_ of node *i* where 0 represents the root.

The rest of the move follows analogously to the *add or remove node* move, including the weighted version.

#### Genotype-preserving prune and reattach

Finally we introduce a variant of *prune and reattach* where instead of retaining the event vector at the pruned node, we retain its genotype of the full copy number profile at that attachment point. When the node is reattached, the event vector of the pruned node is simply updated as the difference in the previous genotype of the pruned node and that of the new attachment point. In the example of Figure A7a the total genotype at event node 2 is (+*S*_1_ + 2*S*_2_ + *S*_3_) while if we were to reattach below event node 5 with genotype (+*S*_1_ + 2*S*_2_) we would simply replace the event vector at event node 2 by +*S*_3_. As the copy number states at each event node are preserved, the likelihood of the current and proposed tree are identical and only the proposed event vector prior needs to be computed and compared to the current value to decide on the acceptance of the move.

Since computing the score of a proposal just involves evaluating the event prior, for this move we score the entire neighbourhood of prune and reattach proposals (including the current tree), and sample directly from the neighbourhood.

#### Changing the overdispersion

Alongside tree moves, we also change the overdispersion parameter *ν*. Since this parameter is positive we work on the log space and propose a new value *ν*′

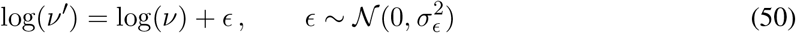

using a Gaussian random walk. The standard deviation of the walk can be adapted to the standard deviation of the recent log(*ν*) values of the chain [with a minimum]. The move to *ν*′ is accepted with probability

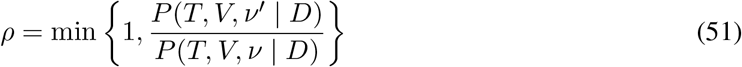

since the move is symmetric.

#### Adaptive tempering

To speed up the search procedure for the highest scoring tree, we can raise the score to a power *P* (*T, V* |*D*)^*γ*^, and adjust the likelihood landscape by varying *γ*. For *γ* > 1 we amplify the differences between tree scores and spend more time exploring a local neighbourhood, while for *γ* < 1 we flatten the landscape and move globally more easily. We vary *γ* adaptively by keeping track of the acceptance probability of moves in the chain. For every tree move which is accepted we transform *γ* → *γ*e^*a*(1−*a*)^ and for every move that is rejected 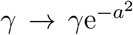 so that we aim to keep the acceptance probability at *a*. As a default we set 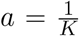 as with more segments it is less likely to pick one that improves the tree.

#### Maximisation instead of summation

Instead of the summation of Equation (9), we can target the score defined by attaching cells at the maximally scoring placement using Equation (10) which avoids the need to penalise the combinatorial complexity. In the scheme above, we simply replace *P* (*T, V* |*D*) everywhere by *S*(*T, V* |*D*) and search for the maximal score *S*. Since we no longer need to satisfy detailed balance, we simplify the acceptance probability of each move to

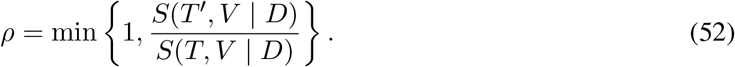

#### Extract common ancestor

In the maximisation mode, we additionally propose to select a pair of sibling nodes uniformly at random, identify the set of common events between them, remove those from each node in the pair, and add them in a new node that is inserted as their new parent. This does not affect the genotypes of the pair of nodes, but may provide a more parsimonious event history. Since the move is not reversible on its own, we preclude it from the sampling scheme and employ it for maximisation.

#### Expand or shrink event

In the maximisation mode, we also include an extension of the *add or remove events* move in which we expand or shrink existing event blocks. We first select a node uniformly at random. Then we choose an event block of a node selected uniformly at random, select either its start or end, and then either remove the event at the edge or expand it to the next contiguous region. Since expanding blocks may merge with others, the reversibility of the move could be quite involved and for simplicity we employ this move only for maximisation.

### E Detailed comparison with CONET

In the main simulation results (Figure 5), CONET performs poorly compared to all other methods, and worse than the results reported in their paper [35]. Indeed, when we run CONET on the simulated data with 2X read depth per bin, it typically finds a large tree with each bin having its own events and does not find the true copy number signal. At these very low coverages, CONET does not have much information from the breakpoint count differences which it relies on for its likelihood computations and tree inference. To recreate results similar to those reported for CONET, we consider their simulation scheme whereby normalised single-cell whole-genome sequencing read depths were generated from copy number profiles by adding Gaussian noise with copy-number-specific variance. In their low-noise setting, these variances are (0.2, 0.01, 0.03, 0.01, 0.07) for copy number values of (0, 1, 2, 3, 4), respectively, and they are doubled for the high-noise case. CONET also assumes a Gaussian noise model for the corrected read counts in the inference.

Since read count data is naturally discrete, this Gaussian approximation corresponds to rescaling discrete data to a mean level of 2 for diploid cells. Since a Gaussian model can also correspond to having no overdispersion, we first compute the effective coverage needed for a discrete model to match the noise in their simulations [35]. For a diploid cell, the coefficient of variation (ratio between standard deviation and mean) is 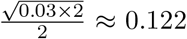 in the high-noise setting, and to get the same variation from a Binomial distribution (as the marginal distribution of each bin) would require

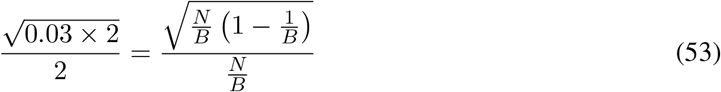

where *N* is the total number of reads, *B* the number of bins with the probability of a read landing in the bin being 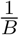, and 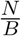 is the per-bin coverage. Solving for the coverage (assuming a large number of bins) we find

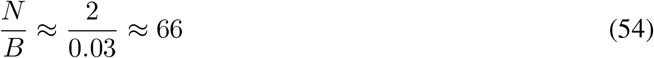

so that the simulation setting of [35] corresponds to coverage of around 66X for their high-noise case and twice as high in their low-noise setting. Any overdispersion in the data generating process would also correspond to even higher coverage.

Although these coverages are much higher than in our simulation (Figure 5), and read depths of very shallow single-cell whole genome sequencing as in [18], they are comparable to that of the datasets of [16, 17]. We therefore repeat our simulations at higher coverages and without overdispersion (Supplementary Figure A5). Although CONET starts to improve at the higher coverages, and gives results closer to their reported ones (bottom row of Figure 3 of [35]) its performance is still worse than Ginkgo, SCOPE and SCICoNE. SCICoNE even starts to give perfect reconstruction in 10-15% of cases.

Though we can come closer to CONET’s previously reported performance by upping the coverage, the authors of CONET [35] claim to outperform SCICoNE which is not the case in our simulations. One potential source of errors is that SCICoNE works directly with the read count data, and assumes a Dirichlet-multinomial distribution of the read counts per bins which allows for some overdispersion. However, the authors of [35] gave to SCICoNE not the actual read counts per bin, but normalised versions rescaled to 2 for diploid regions. When this data is used as input to SCICoNE, it will assume the average per-bin read depth is therefore around 2, and the likelihood component of SCICoNE will be artificially damped down by a large factor.

To illustrate this, we generate data according to the Gaussian simulation model of CONET [35], and use their code for doing so. At these high coverages, we can detect breakpoints without a sliding window and set the window size to a single bin.

For regions with copy number state 0, for which we might expect no reads, the default standard deviation of around 0.45 (variance of 0.2) produces a lot of reads, so we also consider a lower standard deviation of 0.045. As a default we use a very low *η* = 10^−4^ to model regions with no copies so that reads in those regions should be unlikely. To make the comparison more sensible, we set the value *η* for SCICoNE to 0.4 for both standard deviations. When SCICoNE is given as input raw read-count data, we see a distinct improvement in the performance of SCICoNE over CONET (Supplementary Figure A10a). However, when SCICoNE is given the normalised read counts as input instead of the raw counts its performance suffers greatly (Supplementary Figure A10a) since the model is being provided with artificially lowered coverage, and treats this input as observed read counts. Additionally, at such low supposed coverage, we require a sliding window to be able to detect breakpoints and set the window size to 20. The artificially worsened results in Supplementary Figure A10a are very much in line with those reported in the comparisons of [35], while the results here show that by simply giving SCICoNE the correct input it outperforms CONET, in line with our results in the main paper. These results extend to the tree distance measure (Supplementary Figure A10b) though here the ‘sum’ version of SCICoNE is closer to CONET, with a wider spread, while the ‘max’ version of SCICoNE clearly offers a distinct improvement.

**Figure A10:**
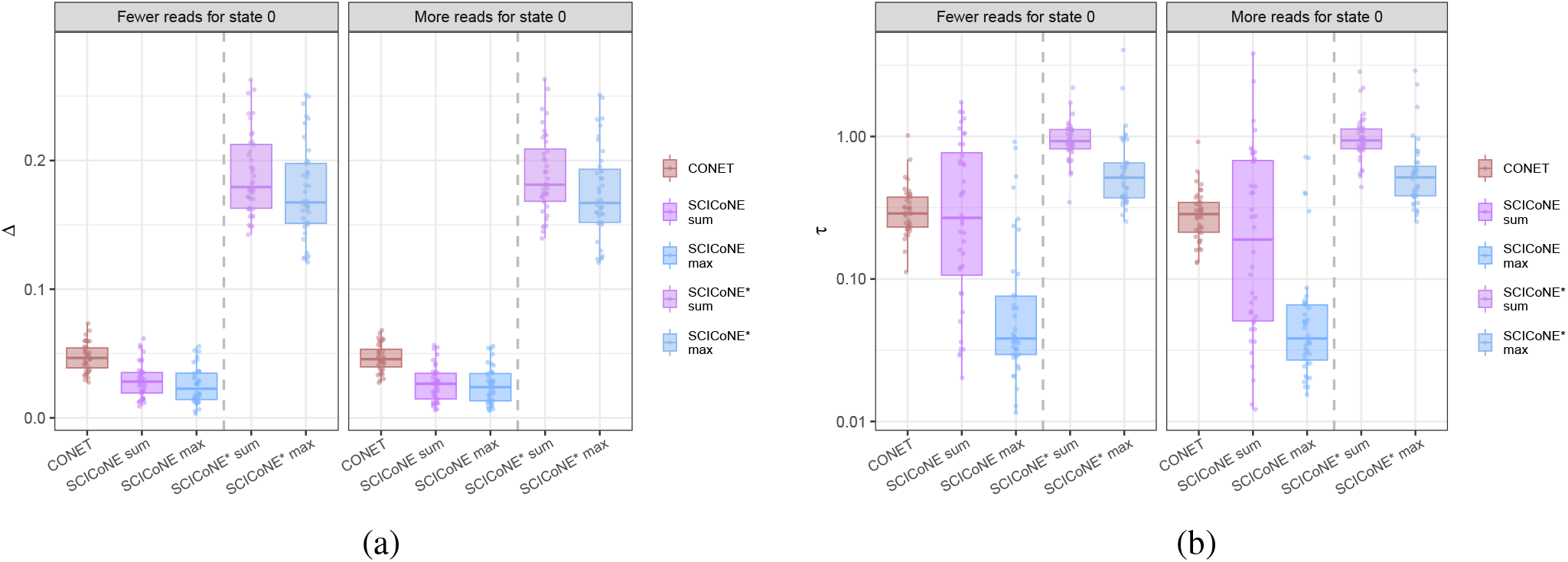
Comparison of copy number calling and tree reconstruction for CONET simulated data. We generate data as in [35] and run both CONET and SCICoNE. Additionally, we run SCICoNE with normalised read data rather than read counts, indicated by the asterisk, with results on the right hand side of the plots. (a) Comparison in terms of copy number calling. (b) Comparison in terms of the tree distance *τ* (Methods) between the true and inferred tree for both CONET and SCICoNE.

These simulations favour CONET since their data simulator matches their modelling and inference scheme. However, we can further disadvantage SCICoNE by using the default value of *η* = 10^−4^ so that we have a large mismatch between the simulated number of reads under the CONET model for regions with no copies and the expected number from this default in SCICoNE. Even with such heavy model misspecification, SCICoNE still outperforms CONET (Supplementary Figure A11), and is easy to improve further (as in Supplementary Figure A10) by simply adjusting *η* to more sensible values for the simulation setting.

**Figure A11:**
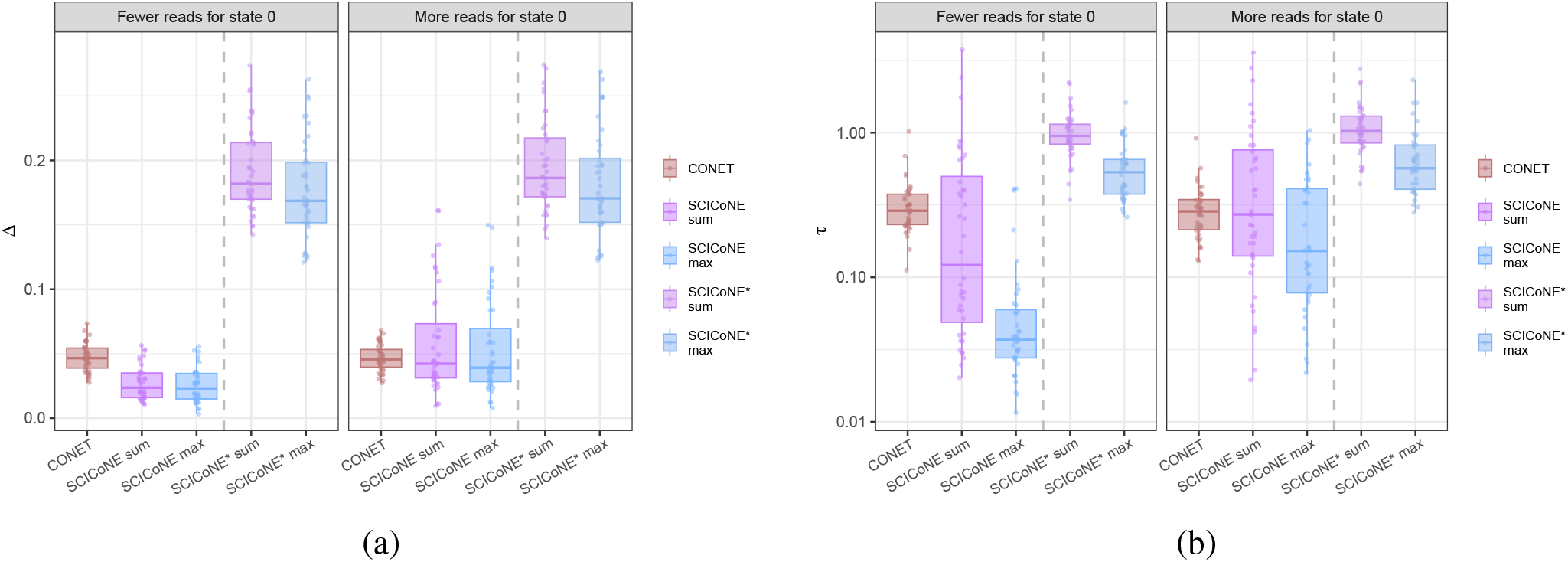
Comparison of copy number calling and tree reconstruction for CONET simulated data. The comparison is as in Supplementary Figure A10 but we use the default SCICoNE model with a very low number of expected reads in regions with no copies and the parameter value *η* = 10^−4^, which is strongly misspecified compared to the data generating process.

